# The inhibitory mechanism of a small protein reveals its role in antimicrobial peptide sensing

**DOI:** 10.1101/2022.12.22.521624

**Authors:** Shan Jiang, Lydia C. Steup, Charlotte Kippnich, Symela Lazaridi, Gabriele Malengo, Thomas Lemmin, Jing Yuan

## Abstract

A large number of small membrane proteins have been discovered in bacteria, but their mechanism of action has remained mostly elusive. Here, we investigate the mechanism of a physiologically important small protein, MgrB, which represses the activity of the sensor kinase PhoQ and is widely distributed among enterobacteria. The PhoQ/PhoP two-component system is a master regulator of the bacterial virulence program and interacts with MgrB to modulate bacterial virulence, fitness, and drug resistance. A combination of crosslinking approaches with functional assays and protein dynamic simulations revealed structural rearrangements due to interactions between MgrB and PhoQ near the membrane/periplasm interface and along the transmembrane helices. These interactions induce the movement of the PhoQ catalytic domain and the repression of its activity. Without MgrB, PhoQ appears to be much less sensitive to antimicrobial peptides, including the commonly used C18G. In the presence of MgrB, C18G promotes MgrB to dissociate from PhoQ, thus activating PhoQ via derepression. Our findings reveal the inhibitory mechanism of the small protein MgrB and uncover its importance in antimicrobial peptide sensing.

**Significance Statement:** Small proteins have high prevalence, vast diversity, and primarily regulatory functions in biological processes across all domains of life. However, their mechanisms of action remain largely elusive. In this study, we investigate the mechanism of the small protein, MgrB. It interacts with the sensor kinase PhoQ, rearranges its conformation, represses its kinase activity, and regulates bacterial response to environmental changes. In particular for antimicrobial peptides, MgrB is required for bacteria to have a selective response to this host-exclusive stimulus. Our findings underline the importance of a small protein in bacterial fitness and drug resistance and provide a molecular basis for engineering novel peptide-based regulators.

## Introduction

Small proteins are defined by their short sequences (usually < 50 aa) and their direct synthesis via ribosomal translation. An increasing number of small proteins have been identified in individual species in all domains of life and complex biological communities, suggesting a high prevalence and vast diversity in biological systems (reviewed in [1–4]). Many of these small proteins (∼30%) are located in the cell membrane or have been predicted to be located there (e.g., [5, 6]). Although only a few detailed studies of small membrane proteins have been conducted, results show that small proteins may target larger proteins and the lipid bilayer, regulating protein function, abundance, and changing membrane properties, demonstrating their predominant role as regulators in the lipid bilayer (reviewed in [7]).

The membrane located PhoQ sensor kinase, a master regulator of a bacterial virulence program, detects host-associated stimuli (review in [8]), such as cationic antimicrobial peptides (CAMPs) [9], mild acidic pH [10, 11], increased osmolarity [12], biliary unsaturated long-chain fatty acids [13], and magnesium limitation [14]. Three small membrane proteins have been identified to control PhoQ activity via direct protein-protein interactions. In *E. coli,* MgrB (47aa) and SafA (65aa) modulate PhoQ activity, with MgrB acting as an inhibitor and SafA as an activator [15, 16]. Deletion of *mgrB* or *safA* has physiological consequences, resulting in a cell division block under magnesium starvation or lowered acid resistance, respectively [17, 18]. In *Salmonella enterica* serovar Typhimurium, a species-specific small protein, UgtS (34 aa), was discovered recently and shown to control the timing of virulence gene expression by preventing premature PhoQ activation when inside macrophages [19].

The PhoQ sensor kinase is found as a dimer on the bacterial inner membrane. It consists of a periplasmic, a transmembrane (TM), a HAMP, a DHp, and a catalytic-ATP binding domain. In the periplasmic domain, several molecular interactions are essential for the functional state of the PhoQ molecule. These include the central hydrogen bonding network (T48 D179 K186) in the PAS-fold [20–23], the inter-monomeric salt bridge (D179-R50’) [24], the membrane-facing acidic cluster (^148^EDDDDAE) and the juxta-membrane region at the periplasm/inner-membrane interface [9, 23, 25]. Mutations altering these molecular interactions change the conformation of the periplasmic domain and the sensitivity of PhoQ to magnesium, low pH, and/or CAMPs. Additionally, different stimuli may result in distinct conformational states of the periplasmic domain, at least in an *in vitro* setting. For example, the PhoQ periplasmic domain showed divergent NMR spectra in low pH compared to that in low magnesium conditions [10]. Besides the periplasmic domain, the PhoQ transmembrane domain was shown to detect environmental osmotic upshifts, where the transmembrane four-helix bundle adopts a more compact and kinase-active conformation [12].

Among the known PhoQ-regulating small proteins, MgrB is the most widely distributed PhoQ repressor in enterobacteria. In *Salmonella*, the deletion of *mgrB* was shown to affect its infection of macrophages and epithelial cells [26]. In *Klebsiella pneumonia*, the disruption of the *mgrB* gene is the most prevalent among other chromosomal mutations in colistin-resistant strains [27]. The expression of *mgrB* is PhoPQ dependent. Thus, the inhibitory function of MgrB provides negative feedback to the two-component system resulting in a partial adaptation of *E. coli* to environmental changes [15, 28]. MgrB is a bitopic inner membrane protein with a short cytosolic N-terminus and a periplasmic C-terminal region (Fig. 1A). Two conserved cysteines (C28 and C39) in the periplasmic region were shown to be essential for MgrB function and suggested to form a disulfide bond [15]. This disulfide bond was proposed to allow MgrB to act as a sensor for the redox state of the bacterial periplasm and create an entry point for redox sensing by the PhoQ/PhoP system [29]. Besides the two cysteines, eight functionally essential residues were identified spread across the entire molecule [30]. However, the mechanism of MgrB inhibiting PhoQ remained largely unclear.

**Fig. 1.**
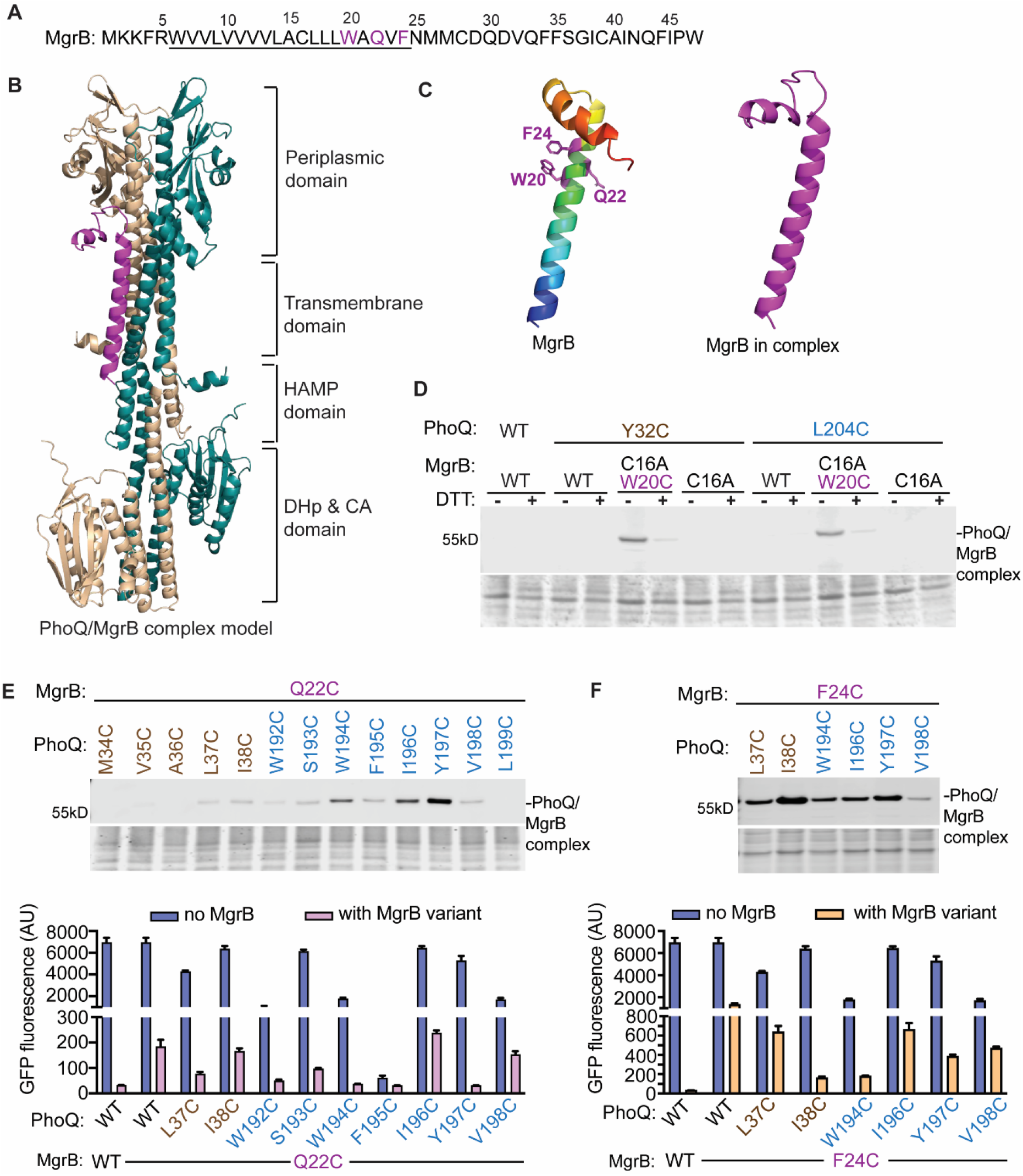
Identification of the interacting residues between MgrB and PhoQ in the transmembrane helices. (**A**) The sequence of MgrB from *E. coli*. Residues predicted to be in the transmembrane helix are underlined. (**B**) The tertiary structure of the top ranked PhoQ/MgrB complex predicted by Alphafold2. The two PhoQ protomers are colored in wheat and teal. MgrB is colored in magenta. (**C**) The structural models of MgrB predicted by Alphafold2 are shown in ribbon. The model of MgrB alone (left) is colored in rainbow spectrum with the N-terminus in blue and the C-terminus in red. The positions involved in crosslinking (20, 22 and 24) are in purple and shown with side chains in ball-and-stick. The predicted structure of MgrB in complex (right) is colored in magenta. All structural figures were prepared using PyMOL unless otherwise stated. (**D, E top, F top**) Western blot analysis of *E. coli* membrane extracts after disulfide crosslinking. The indicated PhoQ and FLAG-tagged MgrB variants were expressed in *E. coli*, followed by Cu-phenanthroline catalyzed disulfide crosslinking (details in Materials and Methods). The total protein stain of the PVDF membranes serves as loading controls. Residues from PhoQ TM1 and TM2 are colored in wheat and teal, respectively. Magenta indicates the residues in the MgrB TM helix. Data are representative of at least three independent experiments**. (D bottom, F bottom**) The functional assay of PhoQ Cys-variants with or without MgrB. An GFP reporter plasmid carrying the *gfp* gene fused with a PhoP regulated promoter P*_mgtLA_* was introduced to the *E. coli* strain. The expression of *phoQ* and *mgrB* variants was induced and cells were grown to the early log phase. The function of PhoQ variants was then monitored by measuring the GFP fluorescence of the cell culture. Each data point is shown with the calculated average and standard deviation from three independent biological replicates

In this study, we combined crosslinking approaches with functional assays and protein dynamic simulations to identify interactions between PhoQ and MgrB, probe MgrB-induced PhoQ conformational changes, and examine the impact of MgrB on PhoQ sensing environmental stimuli. We propose that the binding of MgrB (i) changes the conformation of the linker region between PhoQ periplasmic and transmembrane domains, (ii) rearranges the transmembrane four-helix bundle, and (iii) initiates the translocation of the catalytic domain. MgrB interferes with PhoQ signaling, resulting in an overall lowered activity but still allows PhoQ activation under low pH, osmotic upshift, and magnesium limitation. We further reveal that MgrB mediates PhoQ sensing antimicrobial peptides, enabling bacteria to have a selective response to this exclusively host-associated stimulus.

## Results

### Structural predictions by Alphafold2

We used Alphafold2 multimer [31–33] to predict the structures of the PhoQ/MgrB complex as well as the individual component of the complex (Fig. 1B, C, and S1). The overall prediction of structural elements agrees with previous experimental evidence. For example, the two conserved periplasmic cysteine residues (C28 and C39) in MgrB were positioned in close proximity in the models, structurally supporting the presence of a disulfide bond. In the PhoQ predictions, the PAS fold in the periplasmic domain, the HAMP, the DHp, and the cytosolic catalytic-ATP binding domains resembled the experimental results of these structural motifs [24, 34]. Notably, the region connecting the periplasmic and TM domains and that connecting the HAMP and DHp domains showed a lower predicted local distance difference test loss (pLDDT) score (colored in yellow to green in Fig. S1), indicating these linker regions might have a higher capability of adopting different conformations. Unlike the asymmetric crystal structure of the PhoQ sensor domain [24], the Alphafold2 predicted PhoQ dimer was symmetric, which likely represented only one of many conformations that PhoQ could adopt. In all five PhoQ/MgrB complex models (Fig. 1B and S1), MgrB was placed in between the two PhoQ protomers, with the transmembrane helix parallel to the PhoQ TM helices and the C-terminal short helix (A40- W47) protruding away from PhoQ. The five predicted complex models had only minor differences regarding the positioning of MgrB. Because the PhoQ dimer was mostly symmetrical, Alphafold2 randomly placed MgrB on one or the other side of the PhoQ dimer (Fig. S1). When bound to PhoQ, MgrB showed a major conformational change in the region flanked by the two conserved cysteines (C28 and C39). This region had reduced helicity and rigidness, appearing as an extended flexible loop (Fig. 1C).

### Disulfide crosslinking between MgrB and PhoQ TM helices

Before Alphafold was available, we used the disulfide crosslinking approach to identify the interacting residues in PhoQ and MgrB. We started with residue W20 in the MgrB TM helix, which was suggested to have direct contact with PhoQ [30]. Sequence alignment of MgrB homologs from different bacterial species revealed both W and Y conserved at position 20 (Fig. S2). The MgrB W20Y variant also showed strong inhibitory activity comparable to the wild type (Fig. S2). Since both W and Y have an aromatic ring structure and a polar moiety, we predicted that the nature of the interactions at position 20 likely include hydrophobic interactions and hydrogen bonding. Scanning through the sequence of the two PhoQ transmembrane helices, Y32 in TM1 appeared to be the optimal candidate to fulfill both types of interaction with MgrB W20.

To test this prediction with disulfide crosslinking, we constructed a PhoQ TM_Cys library and generated the MgrB C16A W20C variant, which has only one cysteine residue in the transmembrane helix. The Cys to Ala mutation at MgrB position 16 did not appear to have a functional effect (Fig. S2B). However, the Trp to Cys mutation at position 20 significantly reduced MgrB activity (Fig. S2A). To compensate for the loss of function, we increased the induction of MgrB C16A W20C (pTrc promoter with 10 µM IPTG), which ensured significant repression of PhoQ during crosslinking (Fig. S2D). Additionally, a FLAG tag was fused to the N-terminus of MgrB (Fig. S2E) to facilitate the detection of the PhoQ/MgrB complex in Western blot. Cu-phenanthroline was added to the cells to catalyze the disulfide formation between sulfhydryl groups at a close distance. When MgrB C16A W20C was co-expressed with PhoQ Y32C, we observed a higher molecular weight band at around 55 kDa on the Western blot corresponding to the PhoQ/MgrB complex (Fig. 1D, S3A). This band was reduced by the addition of dithiothreitol (DTT) and was absent when the cysteine residue was at position 16 (WT), or no cysteine was in the MgrB transmembrane helix (C16A). The results are consistent with our prediction suggesting the close proximity of MgrB W20 with the PhoQ TM1 residue Y32 in the complex. Additionally, we found that PhoQ L204C crosslinked with MgrB W20C, indicating position 20 near L204 in PhoQ TM2 as well (Fig. 1D, S3A). Furthermore, we tested PhoQ Cys variants in the neighboring positions of 32 and 204 (PhoQ 31C, 33C, 203C, and 205C). None of the variants formed detectable crosslinked complexes with MgrB W20C in the Western blot (Fig. S2G). Functionally, both PhoQ Y32C and L204C mutants partially restored the MgrB inhibition (Fig. S2F), supporting the hypothesis that MgrB W20 interacts with residue Y32 in PhoQ TM1 and L204 in PhoQ TM2.

Using MgrB residue 20 as a reference point, we next identified interacting amino acids of two more functionally important residues (Q22 and F24) in the MgrB TM helix [30]. In both cases, we observed stronger crosslinking bands and more than one crosslinked position in both transmembrane helices of PhoQ (Fig. 1E, F, S3B, C). For MgrB Q22C, stronger crosslinks were observed to residues in the PhoQ TM2 helix (colored blue). The strongest crosslinking band was shown when PhoQ Y197C was co-expressed. The crosslinking efficiency decreased when the cysteine residue moved to the neighboring positions and further away (196C, 195C, and 198C). It was then partially recovered when the cysteine (194C) was roughly one helical turn away from Y197 (Fig. 1E). Functionally, the Y197C mutation in PhoQ restored the MgrB Q22C activity the best when compared to other PhoQ TM_Cys mutants, supporting the notion that MgrB Q22 primarily interacts with PhoQ Y197.

For MgrB F24C (Fig. 1F, S3C), the strongest crosslinking was found with a PhoQ TM1 mutant, I38C. Functionally, PhoQ I38C showed the highest extent of activity reduction when co-expressed with MgrB F24C, thus restoring the activity of MgrB F24C the best among the tested PhoQ/MgrB pairs. The minor crosslinking events observed could be due to the greater distance between the cysteine residues or an indication of intermediate binding states of MgrB. Notably, one PhoQ Cys variant, F195C, retained only about 1% of the wild-type activity (Fig. 1E). Mutants with such a severe functional defect may have altered MgrB binding, thus unsuitable for the crosslinking approach. For other PhoQ variants in Fig. 1E and F, some showed moderately reduced activity, possibly due to the reduced protein expression level (i.e., PhoQ W192C in Fig. S3F). More importantly, the presence of MgrB significantly repressed the activity of these PhoQ variants, indicating that the change caused by cysteine substitutions does not have a major impact on MgrB inhibition.

Collectively, we identified four-pair residues of MgrB and PhoQ located in close proximity at the protein interface of the complex: W20-Y32, W20-L204, Q22-Y197, and F24-I38. In the AlphaFold2 predicted complex models, these crosslinked pairs were positioned less than 14 Å in the top four ranked models (Table 1). Furthermore, Q22-Y197 and F24-I38 had shorter Cɑ - Cɑ distances than the W20-Y32 and W20-L204 pairs, consistent with our experimental results that the former two pairs showed higher crosslinking efficiency.

**Table 1.**
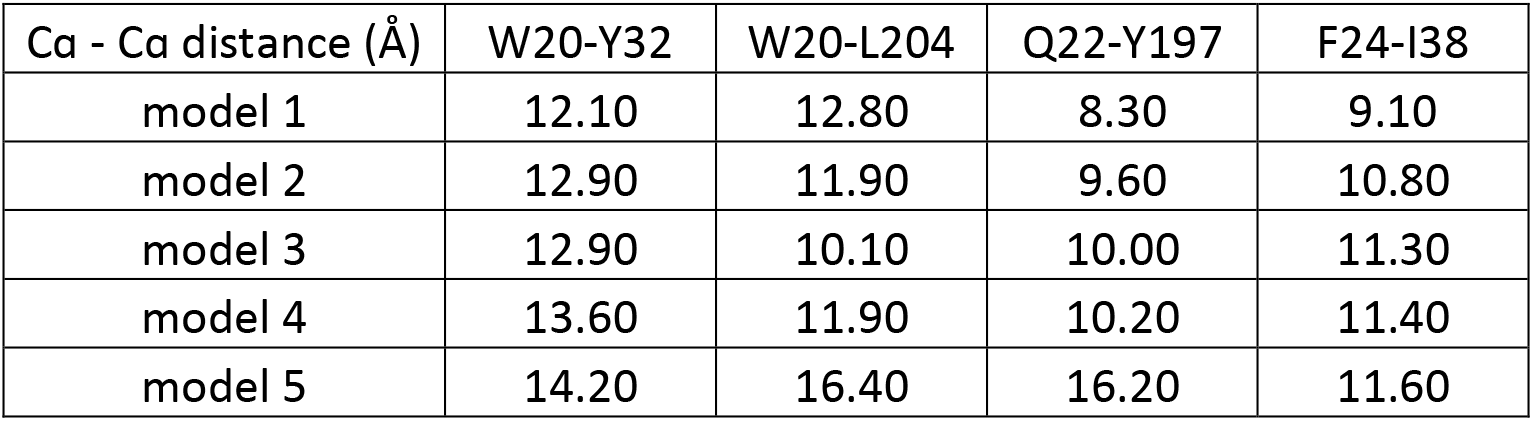
The Cɑ - Cɑ distance of crosslinked residue pairs in the PhoQ/MgrB structural model.

| Ca - Ca distance (Å) | W20-Y32 | W20-L204 | Q22-Y197 | F24-I38 |
| --- | --- | --- | --- | --- |
| model 1 | 12.10 | 12.80 | 8.30 | 9.10 |
| model 2 | 12.90 | 11.90 | 9.60 | 10.80 |
| model 3 | 12.90 | 10.10 | 10.00 | 11.30 |
| model 4 | 13.60 | 11.90 | 10.20 | 11.40 |
| model 5 | 14.20 | 16.40 | 16.20 | 11.60 |

### Crosslinking in the periplasmic domains of PhoQ and MgrB

To gain insights into PhoQ/MgrB interactions at the periplasmic domains, we attempted to identify interacting residue pairs using site-specific photo-crosslinking [35]. We introduced an amber stop codon at each residue in the MgrB periplasmic region (residues 25 to 47). The photo-crosslinking amino acid, *p*-benzoyl-l-phenylalanine (*p*Bpa), was then incorporated into the flag-tagged MgrB at the amber stop codon via the orthogonal tRNA^Amber^ and the aminoacyl-tRNA synthetase in *E. coli*. After UV irradiation, we detected bands with a molecular weight of ∼55 kDa by anti-FLAG antibody when *p*Bpa was incorporated at positions 26, 30, 32, and 38, indicating the formation of the UV-dependent crosslinked PhoQ/MgrB complex (Fig. 2A, S4).

**Fig. 2.**
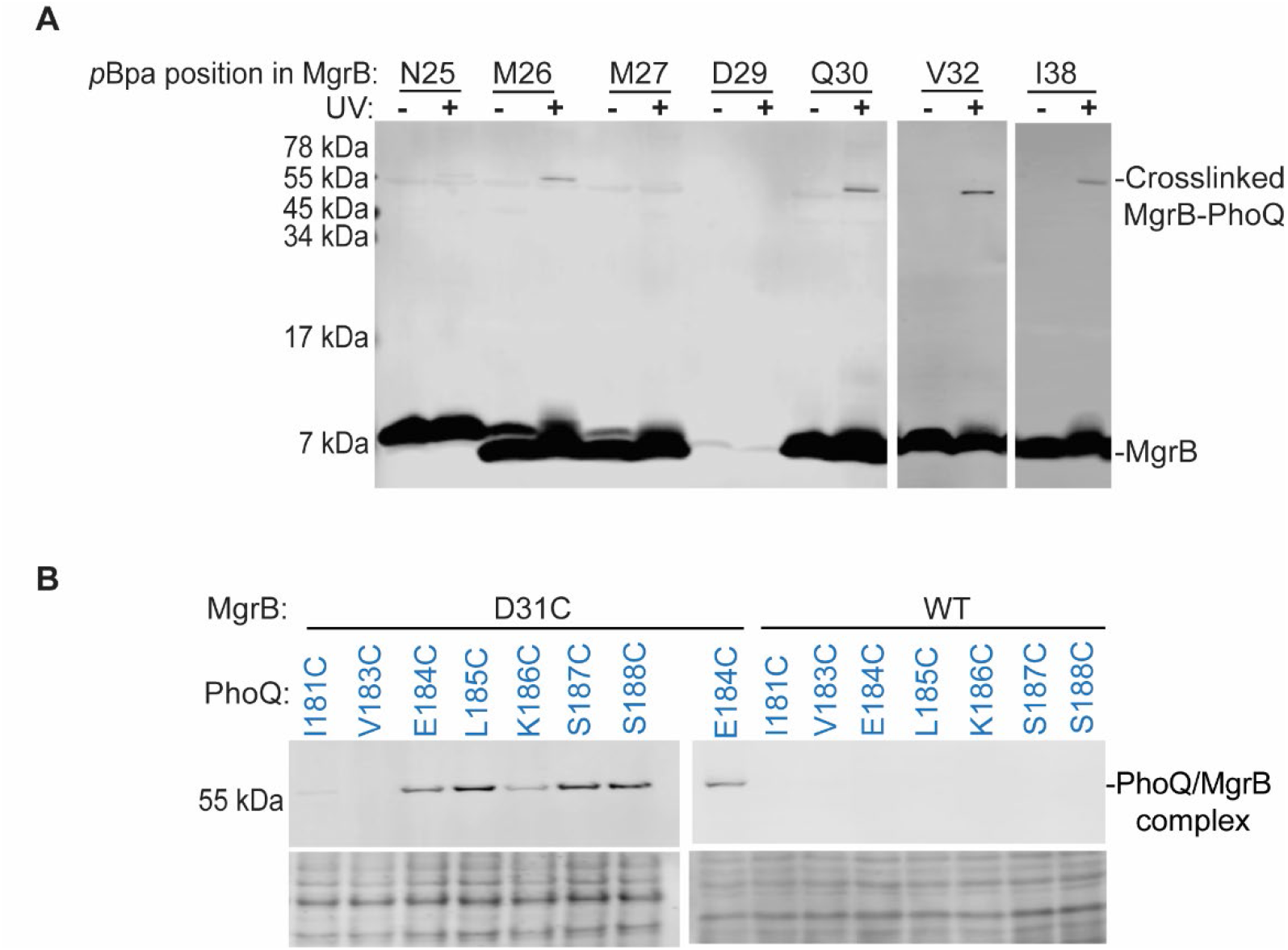
Site-specific photo crosslinking in the periplasmic domain. (**A**) Western blot analysis of the site-specific photo crosslinking between PhoQ and MgrB. The *E. coli ΔphoQΔmgrB* strain harboring pETduet-1 *phoQ-his_6_ flag-mgrB* and pEVOL-pBpF vectors were grown in LB containing the unnatural amino acid, *p*Bpa, which was incorporated into the FLAG tagged MgrB at the indicated positions via the amber stop codon and the cognate tRNA/aminoacyl-tRNA synthetase pair. Cells were treated with UV light. The crosslinked MgrB-PhoQ complex was detected with Western blot using antibodies directed to the FLAG epitopes. Samples without UV irradiation served as controls. (**B**) Western blot analysis of disulfide crosslinking from MgrB residue 31 to the PhoQ linker region connecting the periplasmic domain to the TM2 helix. The experimental procedure is as in Fig. 1D and details can be found in Materials and Methods. Data are representative of at least three independent experiments.

The crosslinked products from positions 30, 32, and 38 could be purified and subjected to mass spectrometry (MS) analysis (Fig. S4 and File S1). PhoQ was detected in all samples as the most prominent protein. Only a small number of other proteins were identified, indicating the protein purification steps were highly effective. MgrB was challenging to detect due to its small size, high hydrophobicity, and a low number of potential MS-detectable peptides. However, we increased the sample injection amount, which allowed for the successful identification of MgrB in crosslinked complex samples, except when *p*Bpa was incorporated at position 38. This was most likely due to the interference of the digestion step with the incorporated *p*Bpa. Despite the successful identification of both PhoQ and MgrB in the crosslinked samples, confident MS identification of MgrB-PhoQ crosslinked peptides providing clear evidence towards positioning the two proteins in the complex was not possible. The lack of crosslinked products with other *mgrB*_*amber* constructs (Fig. S4) could be due to the low efficiency of *p*Bpa incorporation (i.e., position 29) and/or the obstruction of the bulky unnatural amino acid on the binding of MgrB to PhoQ. Most of the residues that resulted in crosslinked products were within the MgrB C28-C39 loop region, suggesting its possible direct interaction with PhoQ.

In the Alphafold2 complex model, the MgrB C28-C39 loop was close to the PhoQ linker region connecting the periplasmic and the TM domains. To test whether the MgrB C28-C39 loop interacts directly with the linker region, we performed disulfide crosslinking with MgrB D31C and PhoQ_linker_Cys variants. Because of two essential cysteines in the MgrB periplasmic region, we first tested if introducing a third cysteine would affect MgrB function. We observed that the introduction of a third cysteine at position 31 did not reduce MgrB inhibition as much as the single cysteine variant, C39A (Fig. S5), suggesting that the cysteine at position 31 does not severely interfere with the C28-C39 disulfide bond formation. Therefore, MgrB D31C is suitable for crosslinking experiments. The crosslinking results showed that MgrB 31C formed disulfide linkages with cysteines in several positions in the PhoQ linker region (184-188) with a similar amount of crosslinked complex (Fig. 2B, S3D). No crosslinked products were detectable in the control sample, where the wild-type MgrB (C28 and C39) was co-expressed with PhoQ_linker_Cys mutants (Fig. 2B, S3E). Our data thus support that MgrB D31 interacts with the PhoQ linker region. The comparable amount of crosslinked complex at multiple positions is probably due to the flexible nature of the MgrB C28-C39 loop.

### The conformational change of PhoQ linker region upon MgrB binding

After mapping the MgrB binding site on PhoQ, we next sought to detect MgrB-induced PhoQ conformational change by comparing the PhoQ dimer crosslinking pattern with MgrB to the one without MgrB. For the PhoQ linker region, we focused on residues 187 and 188, which showed different conformations in PhoQ monomer and dimer models (Fig. S6A). These two residues were located at the periplasmic end of the TM2 helix and at a similar lateral position as the TM1 residue 43 (Fig. S6B). By monitoring the crosslinking efficiency between residues at positions 43 and 188, we intended to detect local helical movements upon MgrB binding. In the absence of MgrB, we observed that 43C and 188C formed an intramolecular disulfide linkage, manifesting as a faster-migrating band below the PhoQ monomer band (Fig. 3A). DTT treatment of the same sample retarded the migration to the exact location as the PhoQ monomer, excluding the likelihood of protein degradation. Nearly no PhoQ dimer band was detected under this condition, further supporting that the orientation of the 188C side chain was more toward the TM1 helix of the same protomer. In the presence of MgrB, the crosslinking pattern changed. The intramolecular crosslinking band reduced, and the intensity of the PhoQ dimer band increased (Fig. 3A, S6C). In this case, crosslinked PhoQ dimer could be formed in four ways: 43C-43’C, 43C-188’C, 43’C-188C, and 188C-188’C. Our control samples showed that the crosslinking was reduced for 43C-43’C and 188C-188’C when MgrB was present. Therefore, the increase in PhoQ dimer formation was likely due to the inter-monomeric crosslinking between positions 43 and 188, suggesting that the binding of MgrB changes the crosslinking pattern from intra- to inter-monomeric linkage and likely the relative orientation between PhoQ residue 188 to residue 43 (Fig. 3B). We also observed a similar trend of change in crosslinking for the residue 43 and 187 pair but with reduced intramolecular crosslinking efficiency (Fig. S6C). Taken together, we hypothesize that MgrB may promote a rotational or translational movement of one TM2 periplasmic terminus towards the TM1 from the other promoter. Because the stoichiometry of PhoQ and MgrB remains unknown, whether the conformational change detected here occurs in both PhoQ protomers awaits further investigations.

**Fig. 3.**
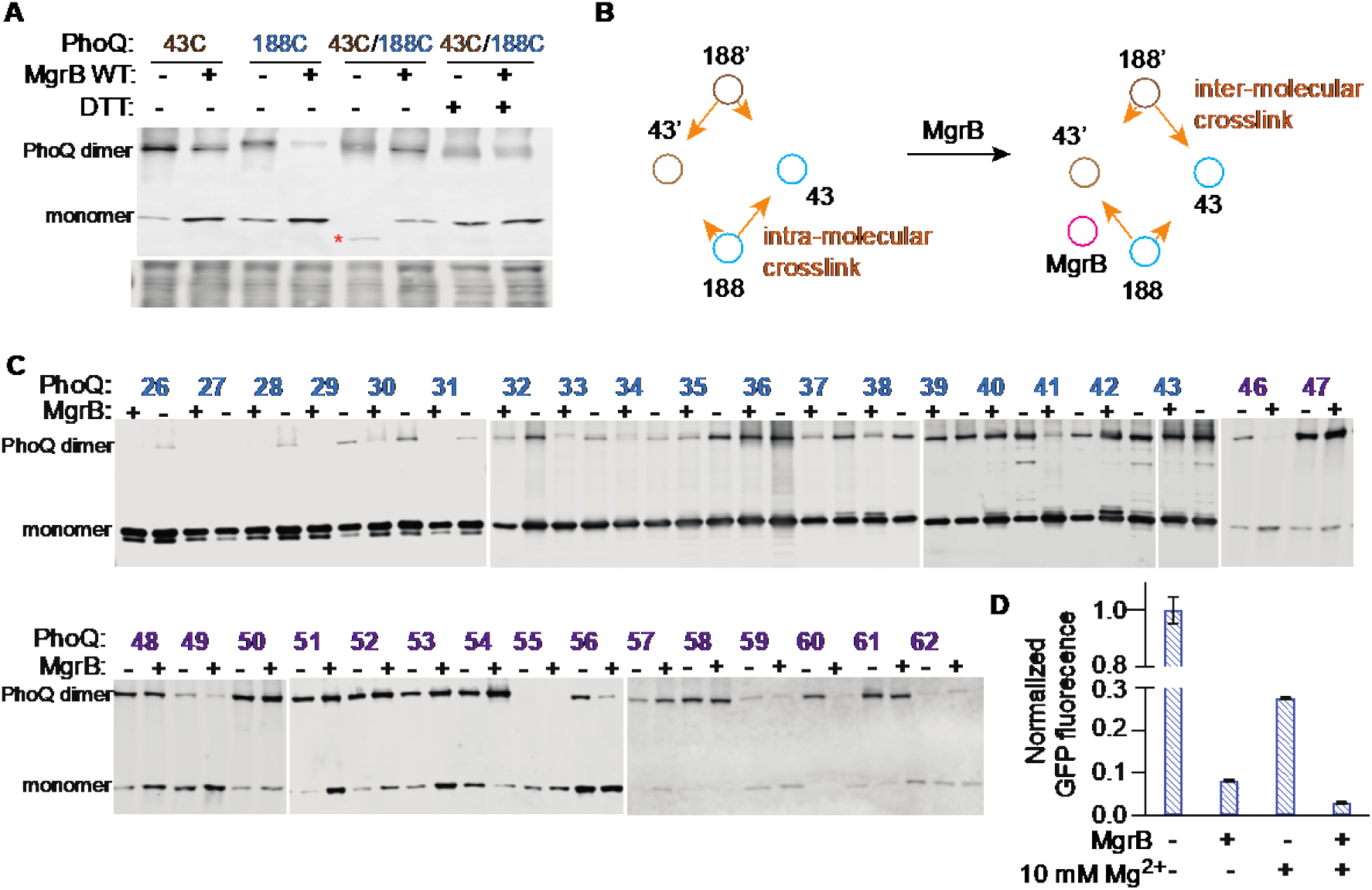
The conformational change of the PhoQ sensor kinase upon MgrB binding. (**A**) Western blot analysis of disulfide crosslinking within the PhoQ molecule in the presence or absence of MgrB. The His_6_-tagged PhoQ cysteine variants were expressed with or without the wild type MgrB in *E. coli*. The disulfide crosslinking reaction was initiated with Cu-phenanthroline. The resulting crosslinked PhoQ species was then detected using Western blot with an anti-His antibody. The red asterisk indicates PhoQ molecules with an intramolecular disulfide crosslink. The total protein stain serves as a loading control. Data are representative of three independent experiments. (**B**) A diagram of MgrB affecting the disulfide crosslinking between the cysteine residues at position 43 and 188. Brown and cyan circles represent the TM helices in each PhoQ monomer. The magenta circle represents MgrB. Orange arrows indicate the crosslinking tendency of Cys188 to Cys43 in the same or different PhoQ monomer. (**C**) MgrB induced PhoQ conformational change along the PhoQ dimer interface. The His_6_-tagged PhoQ cysteine variants were expressed with or without the wild type MgrB in *E. coli*. The experimental procedure is as described in A and details can be found in Materials and Methods. The position of cysteine is indicated as residue numbers in the PhoQ N-terminal helix. Numbers colored in blue and purple are residues in TM and periplasmic domains, respectively. Data are representative of three independent experiments. (**D**) The repression of PhoQ activity by high magnesium and MgrB. The activity of PhoQ was monitored with the GFP reporter system using the *E. coli ΔphoQΔmgrB* strain as described in Fig. 1E. Data are representative of three independent experiments. Each data point is shown with the calculated average and standard deviation from three biological replicates.

### MgrB repression of PhoQ is divergent from high magnesium repression

Besides the small protein MgrB, high magnesium also represses PhoQ kinase activity [14]. The structural change of PhoQ under magnesium repression was studied in detail by monitoring the arrangement of α helices along the PhoQ dimer interface via disulfide crosslinking [36]. It was proposed that under high magnesium conditions, the N-terminal α helix across the TM and the periplasmic domains undergoes a scissoring-type of motion with the pivot point near the periplasm/membrane interface [36].

Using a similar approach, we monitored the arrangement of α helices along the PhoQ dimer interface in the presence and absence of MgrB. Our data showed that most TM1 residues had reduced crosslinking between PhoQ protomers upon MgrB binding (Fig. 3C blue residues, Fig. S7). This reduction was more prominent for residues close to the cytoplasm, suggesting that MgrB binding pushes TM1-TM1’ apart and the propagation of PhoQ conformational change from the periplasm to the cytoplasm. In contrast, the presence of MgrB did not show a similar crosslinking effect on residues along the periplasmic helical interface (Fig. 3C purple residues, Fig. S7). Most residues in this region showed little change in the crosslinking pattern. Some showed a significantly decreased (i.e., residues 46 and 60) or a slightly increased dimer fraction (i.e., residues 62). The overall PhoQ dimer rearrangement upon MgrB binding differed from the scissoring motion proposed for high magnesium repression, suggesting that MgrB and high magnesium act differently on PhoQ. We then did functional analyses of PhoQ under high magnesium, with MgrB, or with both repressive factors. The results showed that PhoQ was further repressed significantly when both high magnesium and MgrB were present (Fig. 3D), supporting the hypothesis that magnesium and MgrB may have divergent mechanisms of repression.

### Molecular dynamics simulations of PhoQ/MgrB complex models

When comparing the predicted PhoQ dimer with its complex with MgrB, there was only minor conformational change, which is in stark contrast to our experimental results. The Alphafold2 prediction might have only captured a transient arrangement of the complex but not the final state. To investigate the effect of MgrB on the conformation of PhoQ in silico, we carried out a set of four molecular dynamic simulations. The initial atomic models were obtained using the best-ranked model predicted by the multimeric version of Alphafold2. In total, we modeled four different complexes: PhoQ dimer (i) alone, and interacting with (ii) the wild-type MgrB, (iii) MgrB W20Y, and (iv) MgrB W20A. Each complex was inserted into a bacterial mimetic membrane bilayer, and simulations were carried out up to 1 μs under near-physiological conditions. To evaluate the impact of the starting conformation on the stability of the PhoQ/MgrB complex, we carried out two additional 500 ns simulations seeded from the second and third predictions. All protein complexes remained stable throughout the entire simulations, with minor fluctuation in the secondary structure. We did not observe any significant differences between the three PhoQ/MrgB simulations. The MgrB C-terminal short helix (A40-W47) was buried close to the membrane surface. The C-terminal carboxylate group interacted with the head group of PE lipids in the simulations. The orientation of this C-terminal helix was primarily restrained by the disulfide linkage (C28-C39) and remained stable in the simulations. A principal component analysis of the transmembrane domain clearly showed that the functional MgrB variants (WT and W20Y) induced a similar conformational change (Fig. 4A), corresponding to an increase in the distance between PhoQ TM1 helices, coupled with a decrease in distance between TM2 helices. This change is more evident in TM residues near the membrane/cytoplasm interface (Fig. 4B, S8). The MgrB W20A had only a marginal impact on the conformation of the transmembrane domain.

**Fig. 4.**
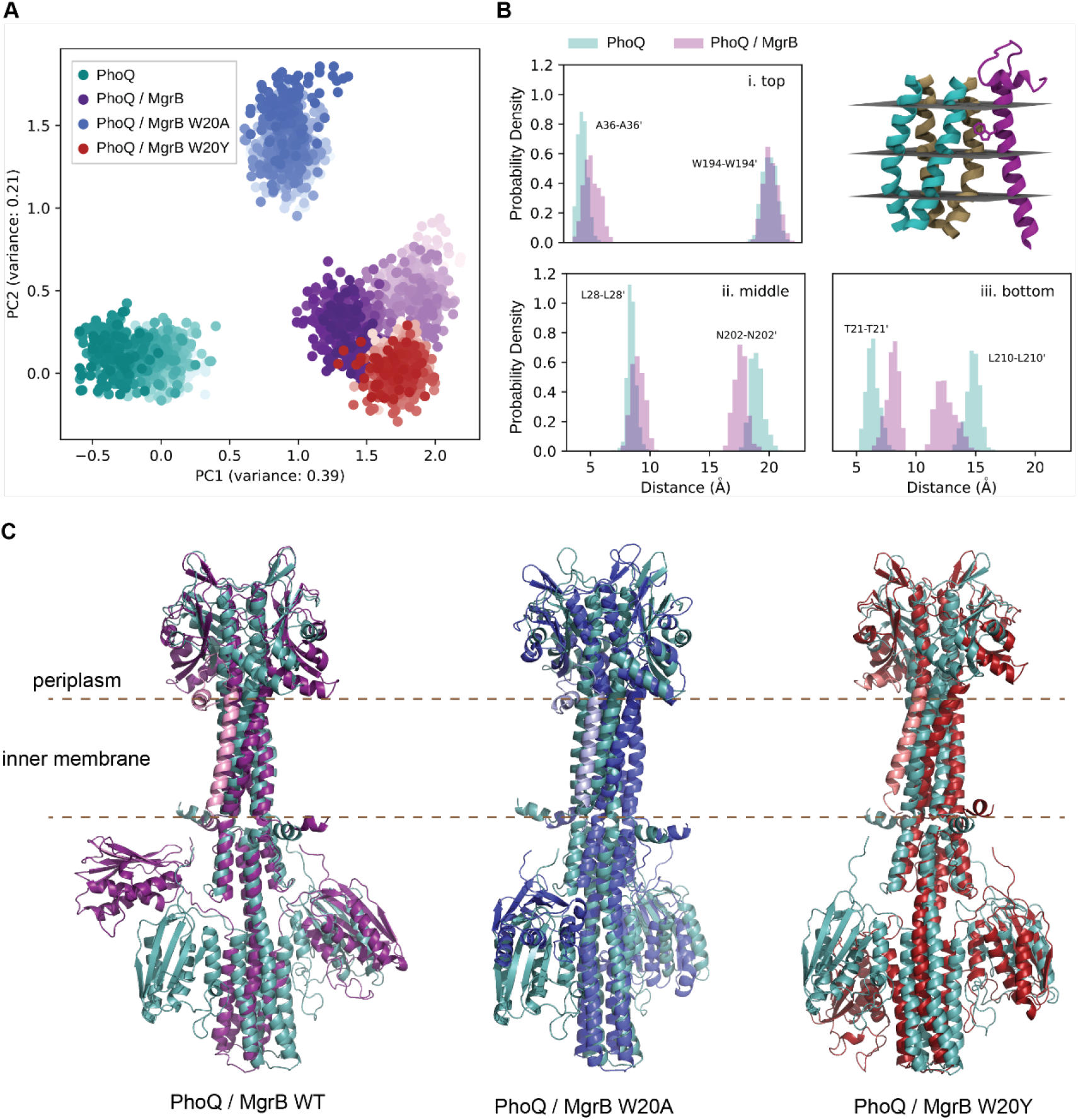
Molecular dynamics simulations of PhoQ/MgrB complex models. (**A**) Projection of the molecular dynamics simulations into the eigenspace formed by the two first components of a Principal Component Analysis (PCA) of the transmembrane domain of PhoQ (residues 21 – 38 and 194 – 215). Each point corresponds to a conformation of the TM and is colored in shades of teal, purple, red and blue, for the simulations of PhoQ alone, and in complex with MgrB, MrgB W20Y and W20A, respectively. Lighter shades correspond to conformations sampled earlier in the simulation. (**B**) Distribution in distances between TM1 – TM1’ (left) and TM2 – TM2’ (right) at the top, middle, and bottom of PhoQ TM domain. (**C**) The overlay of the final conformations of PhoQ/MgrB complex models with PhoQ dimer. PhoQ structures are colored as in A. The MgrB wild-type, W20A, and W20Y variants are colored in pink, light blue, and light red, respectively.

In all the simulations with MgrB variants, we also observed the translocation of one of the PhoQ catalytic-ATP binding (CA) domains (Fig. 4C). This is most prominent in the 1 μs simulations with the wild-type MgrB, where one CA domain moved fully away from the helix harboring the phosphorylatable histidine and located itself close to the inner membrane. For PhoQ alone, we did not observe any significant movement of the CA domains throughout the entire simulation. The functionally impaired MgrB W20A also induced PhoQ CA domain movement, though to a lesser extent compared to the wild type (Fig. 4C). In this simulation, the CA domain remained close to its initial position but rotated by about 90°. Our previous work showed that the W20A mutant had low activity due to reduced affinity to PhoQ [30]. However, since the simulation was seeded with a complex where MgrB W20A was already interacting with PhoQ, this likely represents the scenario where overexpression of MgrB W20A compensates for its weak binding to PhoQ and partially recovers its inhibitory function.

Overall, the simulation results confirm our experimental observations, showing the PhoQ structural changes at the MgrB binding site. Furthermore, MgrB induces movements of the distantly located CA domain, suggesting that the local conformational change due to MgrB binding results in the repression of PhoQ kinase activity.

### MgrB mediates PhoQ sensing cationic antimicrobial peptides

Our crosslinking data (Fig. 2B) suggested that MgrB directly interacts with PhoQ E184, a residue involved in CAMP sensing [9]. We, therefore, hypothesized that MgrB might affect PhoQ activation by CAMPs. To investigate this as well as the impact of MgrB on PhoQ sensing other stimuli, we tested PhoQ activation by cationic antimicrobial peptides, low pH, and high osmolarity in the presence and absence of MgrB (Fig. 5). Under a repressive magnesium condition (LB medium supplemented with 10 mM Mg^2+^), only slight or no PhoQ activation was detected by antimicrobial peptide C18G regardless of MgrB, consistent with previous work where high magnesium repressed the activation of PhoQ by CAMPs in *Salmonella* [9, 10]. Conversely, with the same magnesium condition (10 mM), a relatively large activation of PhoQ was detected by low pH or osmolarity upshift in the presence of MgrB. The activation did not appear to have a qualitative change when MgrB was absent, except that the overall PhoQ activity increased due to the lack of MgrB repression (Fig. 5A).

**Fig. 5.**
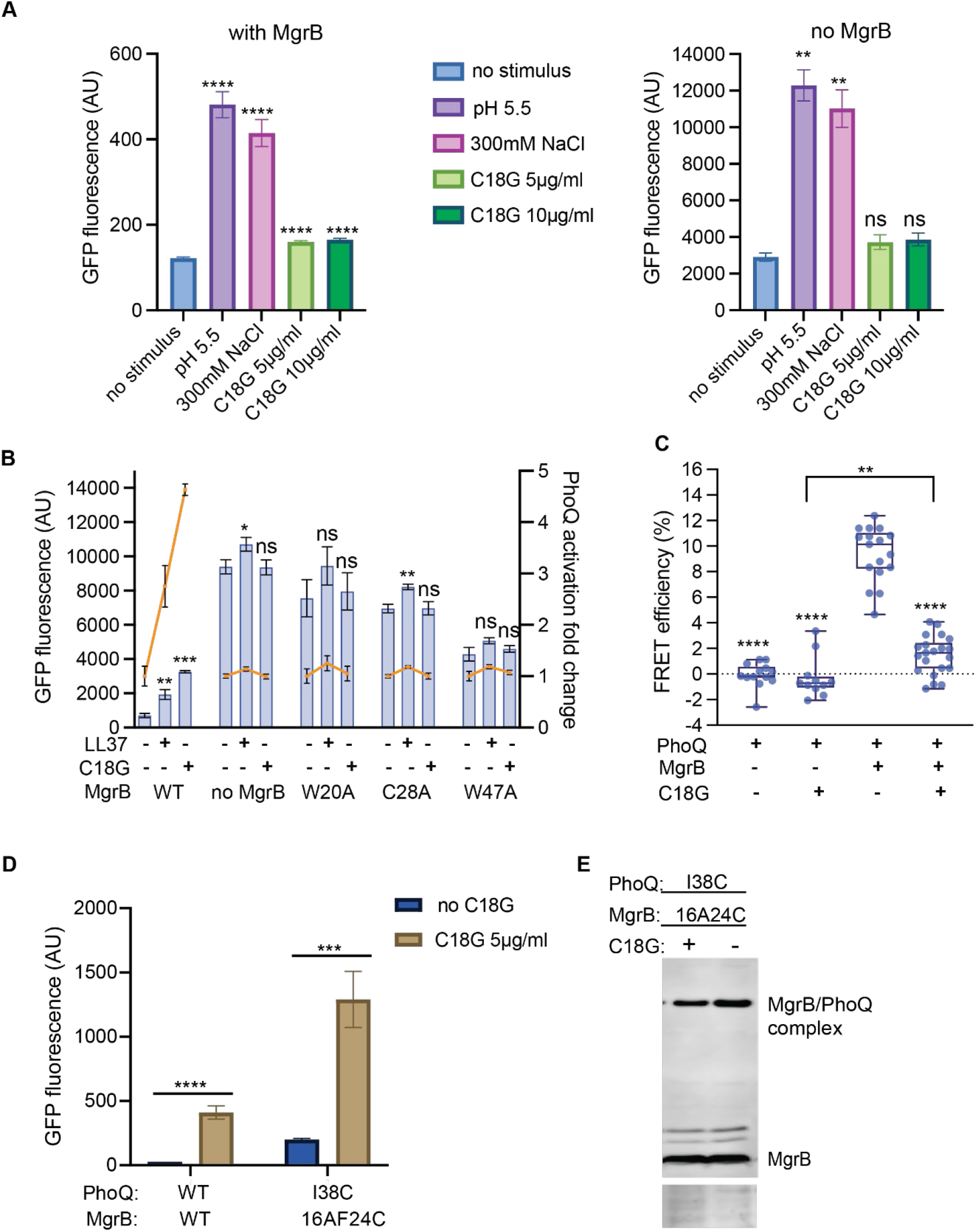
The impact of MgrB on PhoQ sensing external stimuli. (**A**) The PhoQ activation in the presence or absence of MgrB under high magnesium conditions. *E. coli* MG1655 *ΔmgrB* strain harboring pBAD33 with (left panel) or without (right panel) *mgrB* gene, were induced with arabinose and grown in the presence of indicated stimuli in LB media supplemented with 10 mM magnesium. The activity of PhoQ was monitored using the GFP reporter as described in Fig. 1E. (**B**) The activation of PhoQ in the presence of antimicrobial peptides and MgrB under 1 mM magnesium conditions. The orange lines indicate the fold change of PhoQ activity in the presence of antimicrobial peptides. *E. coli* MG1655 *ΔmgrB* strain expressing MgrB variants were grown in LB supplemented with 1 mM magnesium and antimicrobial peptides, LL37 (5 µg/ml) or C18G (5 µg/ml) as indicated. The PhoQ activity was followed as in A. (**C**) The FRET measurements of PhoQ-mNeonGreen and mCherry-MgrB *in vivo*. *E. coli* cells expressing PhoQ-mNeonGreen and mCherry-MgrB from plasmids were grown to mid-log phase, then treated with or without the antimicrobial peptide, C18G (10 µg/ml). The energy transfer from mNeonGreen to mCherry was then measured for each condition. The strains expressing no FRET receptor (mCherry-MgrB) serve as negative controls. (**D**) The activation of PhoQ variants in the presence of MgrB and C18G (5 µg/ml). The growth and induction conditions for the PhoQ and MgrB variants were the same as described before (Fig. 1F bottom). The PhoQ activity was monitored by measuring the GFP reporter fluorescence. (**E**) Western blot analysis of PhoQ I38C MgrB 16A24C disulfide crosslinking in the presence or absence of C18G (5 µg/ml). The crosslinking and Western blot experiments were performed as described before (Fig. 1F top). The total protein stain of the membrane serves as a loading control. Data are representative of three independent experiments. Each data point in A-D is shown with the calculated average and standard deviation from at least three independent experiments.

We next followed PhoQ activities in the presence of antimicrobial peptide LL37 and C18G under a physiologically relevant magnesium concentration (1 mM Mg^2+^ in LB) that does not repress CAMPs from inducing PhoQ activity [9]. Up to a nearly 5-fold increase of PhoQ activity by antimicrobial peptides was detected when the wild-type MgrB was present (Fig. 5B). However, in the absence of MgrB, there was no significant further increase of PhoQ activity in the presence of C18G. In the case of LL37, only a marginal increase (1.14-fold) of PhoQ activity was observed, consistent with a previous report [15], suggesting that LL37 may be slightly different from C18G in terms of PhoQ activation. To test if PhoQ was already maximally activated in the *ΔmgrB* strain with 1 mM magnesium, we treated the cells with additional stimuli. PhoQ activity was significantly further increased by lowered pH or elevated osmolarity (Fig. S9A), indicating that PhoQ activity did not reach saturation with 1 mM magnesium alone. To be precise on the total magnesium concentration in the growth medium, we tested our strains in the defined minimal A medium [37] supplemented with 1 mM magnesium. Consistent with what we observed in LB, PhoQ activity was higher in the absence of MgrB but could not be further increased by C18G. Therefore, we hypothesize that MgrB may mediate PhoQ to sense antimicrobial peptides under a physiologically relevant magnesium concentration.

The lack of PhoQ activation by antimicrobial peptides in the absence of MgrB might also be due to the protective effect of the overall elevated activity of the PhoQ/PhoP two-component system, which could increase the outer membrane modification and reduce the entry of antimicrobial peptides to the periplasm. To obtain more direct evidence, we monitored PhoQ and MgrB interactions *in vivo* using the acceptor photobleaching Förster resonance energy transfer (FRET) approach. Functional PhoQ-mNeonGreen and mCherry-MgrB fusion proteins (Fig. S9C and D) were expressed in *E. coli*. Cells were collected in the mid-log phase, treated with C18G, and then used for FRET measurements (Fig. 5C). The control cells without C18G treatment, showed energy transfer from mNeonGreen to mCherry with about 10% efficiency, comparable to our previous report [12], indicating the formation of the PhoQ/MgrB complex *in vivo*. With the C18G treatment, the FRET efficiency reduced to about 2%, slightly higher than the negative controls, suggesting that C18G induces the dissociation of the PhoQ/MgrB complex.

To confirm the *in vivo* FRET results, we performed disulfide crosslinking experiments with or without C18G using the PhoQ I38C MgrB F24C variant pair. We chose this pair because: i) they formed a good amount of PhoQ/MgrB crosslinked complex readily detectable by Western blot (Fig. 1F); ii) PhoQ I38C showed wild-type-like activity (Fig. 1F); iii) MgrB F24C repressed PhoQ I38C to a relatively low level (Fig. 1F); iv) the variant pair responded to C18G showing significantly increased PhoQ activity (Fig. 5D). Compared to the crosslinking in the absence of C18G, we observed a lighter band corresponding to the PhoQ/MgrB complex (Fig. 5E). Assuming that C18G does not adversely affect Cu: phenanthroline catalysis, the reduction in the crosslinked complex suggests that C18G induces MgrB to move away from PhoQ, consistent with our FRET results.

Taken together, we propose that MgrB is involved in PhoQ sensing CAMPs. Under physiologically relevant magnesium conditions, PhoQ is active but partially repressed due to the bound MgrB. This state allows PhoQ to detect and respond sensitively to CAMPs at a subinhibitory concentration. CAMPs bind to PhoQ and induce the release of the bound MgrB, further activating PhoQ/PhoP-regulated genes and potentially increasing bacterial resistance to CAMPs (Fig. 6).

**Fig. 6.**
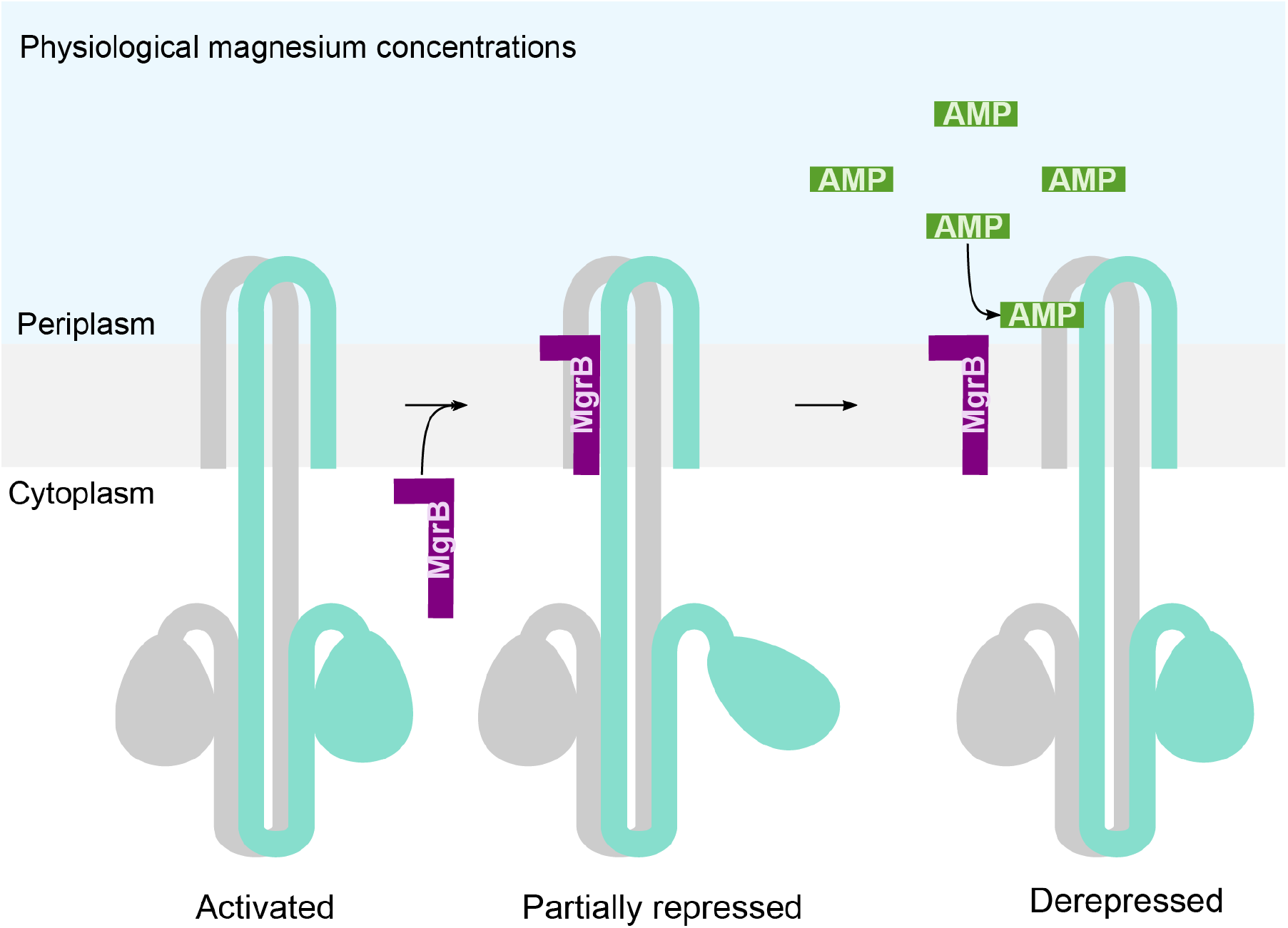
The schematic representation of MgrB-mediated PhoQ activation by antimicrobial peptides. PhoQ is active under physiological magnesium conditions, inducing the expression of *mgrB*. The expressed MgrB binds to PhoQ, resulting in a partially repressed state. Antimicrobial peptides (AMPs) enter bacterial periplasm, bind to PhoQ, and cause MgrB to dissociate from PhoQ.

## Discussion

In this study, we report that MgrB directly interacts with the TM helices and the periplasmic/TM linker region of PhoQ, changing its conformation in a way that divergent from high magnesium repression. It reduces the overall activity of PhoQ yet still allows its activation by low magnesium, low pH, and increased osmolarity. We further reveal that PhoQ activation by CAMPs is MgrB-dependent under physiologically relevant magnesium conditions. CAMPs induce MgrB dissociating from PhoQ and thus increase PhoQ activity via derepression. A higher PhoQ activity leads to an increased outer membrane modification and improved resistance to CAMPs. Since the septum is the initial attacking site of certain CAMPs (i.e., LL37) on the membrane [38], the possible division block caused by strong PhoQ activation may also protect bacteria against the host’s innate immune response. CAMPs were suggested to bind to an acidic surface in the PhoQ periplasmic domain, though the exact binding site has not been well defined [9]. The acidic cluster (^148^EDDDDAE), a major component of the acidic surface, is relatively distant from the MgrB binding site. However, it remains unclear whether the acidic residue E184 in the linker region is directly involved in CAMP binding. It is plausible that the binding of CAMPs induces PhoQ conformational change, which leads to the dissociation of MgrB from PhoQ.

We observed that a high level of magnesium (10 mM) repressed PhoQ activation by CAMPs in the presence or absence of MgrB in *E. coli*, consistent with a previous report on PhoQ in *Salmonella*, though MgrB had not yet been discovered at that time [9]. The same report proposed that CAMPs activate PhoQ by replacing magnesium ions from the acidic cluster in PhoQ, which is different from our MgrB-mediated CAMP sensing mechanism. Both hypotheses, however, do not conflict with each other but rather describe two aspects of PhoQ sensing CAMPs under different magnesium conditions. It is also possible that magnesium may affect MgrB binding to PhoQ, thus connecting these two mechanisms of CAMP sensing by PhoQ. Interestingly, the magnesium replacement mechanism can not explain PhoQ activation by antimicrobial peptides in some cases. For example, a recent study revealed that antimicrobial peptides with a ß-sheet structure or a neutral charge could also activate PhoQ [39]. These newly found PhoQ activators do not possess the properties to interact with the PhoQ acidic cluster. Furthermore, there are PhoQ homologs that do not contain an acidic cluster in their sequences. Our MgrB-dependent antimicrobial peptide sensing mechanism may provide a suitable explanation for both cases.

By monitoring PhoQ conformational change and activities, we show that MgrB repression diverges from high magnesium repression. This is consistent with the different binding sites of magnesium and MgrB on PhoQ. Magnesium was suggested to bind to several locations, including the acidic cluster near the inner membrane (148-152) and residues G93, W97, H120, and T156 [14, 23, 40]. Mutations of these residues were shown to abolish magnesium protection from iron-mediated protein cleavage [40]. The majority of these residues are located in the peripheral of the periplasmic domain, distinct from the MgrB binding site.

Our crosslinking results revealed extensive interactions between MgrB and the PhoQ linker region, implying its importance in functional regulations. Highly conserved and functionally important residues are found in or near this region, such as K186, T48, R50, E184, and L185, mutations of which lead to alterations in PhoQ activity as well as signaling capability [20, 23, 24]. Curiously, the wild-type MgrB maintained its inhibition on PhoQ linker variants, suggesting that the flexible MgrB loop may have various interaction modes with the PhoQ linker region. Interestingly, the PhoQ activating protein, SafA, appears to act near this linker region in terms of the putative SafA binding site, and the activation required inter-monomeric salt bridge (R50’-D179) [41, 42]. Whether MgrB repression competes with SafA activation of PhoQ and how these regulations are coordinated await further studies.

Sensor kinases in bacterial two-component systems are evolutionarily related and have structural resemblance. We found that the predicted structures of EnvZ, QseC, and CreC sensor kinases closely resemble the PhoQ structure. EnvZ was also shown to form a complex with MgrB experimentally [30]. With the mechanism of MgrB revealed here, it is foreseeable that novel small protein regulators can be designed for PhoQ, EnvZ, and possibly many other sensor kinases. This will contribute to the repertoire of new antimicrobial agents targeting sensor kinases that regulate bacterial virulence [43] and to the treatment of infections caused by drug-resistant pathogens.

## Materials and methods

### Bacterial strains, plasmids, and growth conditions

The complete lists of strains, plasmids, and primers used in this study are summarized in Table S1, S2, and S3, respectively. The *E. coli* MG1655 *ΔmgrB* and *ΔphoQΔmgrB* strains were from the previous study [30]. All strains were grown in the lysogeny broth (LB) medium or minimal A medium [37] containing 0.2% glycerol, 0.1% casamino acids supplemented with magnesium as indicated. Cells were grown at 37 °C with shaking unless otherwise indicated. Genes in pTrc99a vector were induced with isopropyl β-D-1-thiogalactopyranoside (IPTG) at a final concentration of 10 μM. Genes in pBAD vector were induced with 0.008% arabinose. Genes in pEVOL-pBpF vector were induced with 0.02% arabinose. Genes in pETduet-1 vector were induced with IPTG at a final concentration of 0.5 mM. When appropriate, the antibiotics ampicillin, kanamycin, and chloramphenicol were used at concentrations of 100 μg/ml, 50 μg/ml, and 34 μg/ml, respectively. A detailed description of cloning and plasmid construction can be found in the SI Materials and Methods.

### Disulfide crosslinking *in vivo*

The *in vivo* disulfide-crosslinking experiments were performed as previously described with modifications [44]. Briefly, cells carrying pBAD33 *phoQ* variant and pTrc99A *mgrB* variant were grown at 37 °C in LB medium supplemented with 10 mM MgSO_4_ overnight, then diluted 1:100 to fresh LB medium supplemented with 1 mM MgSO_4_. The expression of plasmid-encoded proteins was induced with arabinose and IPTG. When the OD_600_ value reached 0.6, cells were harvested, washed, and resuspended in the crosslinking buffer (137 mM NaCl, 2.7 mM KCl, 10 mM Na_2_HPO_4_, 1.8 mM KH_2_PO_4_, 1 mM MgSO_4_, and 0.1 mM Cu-phenanthroline), followed by 10 minutes incubation at 25 °C. The reaction was stopped by pelleting the cells and then resuspending in the 1x stop-solution (8 mM sodium phosphate buffer, pH 7.8, 12.5 mM EDTA, and 12.5 mM N-ethylmaleimide) and incubation at 25 °C for 10 minutes. Cells were collected, resuspended in the lysis buffer (50 mM Tris-HCl, pH 8.0, 100 mM NaCl, and protease inhibitors), mixed with lysing matrix B beads (MP Biomedicals), and broken with a homogenizer following the manufacturer’s instructions. The mixture was spun down at 2,300 x g for 10 minutes at 4 °C to pellet the beads and cell debris. The supernatant was centrifuged at 21,000 x g for one hour at 4 °C to pellet the membrane fraction, which was then dissolved in the SDS sample buffer at 4 °C overnight. The membrane proteins were separated on an SDS tris-tricine or tris-glycine polyacrylamide gel via electrophoresis and analyzed with Western blot.

### Western blot analysis

Proteins were transferred from a polyacrylamide gel to a polyvinylidene fluoride membrane or nitrocellulose membrane. PhoQ-His and Flag-MgrB were detected with the anti-His and anti-Flag primary antibody (Sigma), respectively. The IRDye 800CW-conjugated secondary antibody (LI-COR) was used, and the protein bands were visualized with the Odyssey CLx imaging system (LI-COR) and quantified with ImageJ. The crosslinking efficiency was calculated by dividing the amount of PhoQ/MgrB complex with the total amount of PhoQ/MgrB complex and MgrB monomer.

### Functional assay with the *gfp* reporter gene

The GFP reporter assay was performed as a previous study [12]. Briefly, the reporter plasmid pUA66 *P_mgtLA_-gfp* was transformed to strains indicated. The resulting transformants were grown overnight at 37 °C in LB medium with 10 mM MgSO_4_, then diluted 1:100 to fresh LB medium with magnesium and/or stimuli as indicated. The cultures were grown at 37 °C with vigorous shaking till the early log phase (OD=0.4-0.5). The fluorescence of cells was monitored with a BD LSRFortessa SORP flow cytometer (BD Biosciences) equipped with a 100-mW 488-nm laser and a 510/20 bandpass filter. BD FACSDiva software version 8.0 (BD Biosciences) was used to analyze the acquired data.

### Unnatural amino acid incorporation in MgrB and site-specific UV crosslinking *in vivo*

The site-specific photo-crosslinking experiments were performed as previous studies with modifications [35, 45]. *E. coli* BL21 DE3 cells carrying pETduet-1 *phoQ-his_6_ flag-mgrB* and pEVOL-pBpF vectors were grown in LB medium at 37 °C overnight, then diluted 1:100 to fresh LB medium containing 0.5 mM *p*Bpa. When the OD_600_ value reached around 0.6, cells were induced with arabinose (0.02%) and IPTG (0.5 mM) for 4h at 30 °C, followed by harvesting, washing two times, and resuspension in 1 ml cold PBS buffer. The bacterial suspension was spread out onto a 35 mm round dish on ice and irradiated for 5 minutes with 365 nm UV light using an LED UV hand lamp (Honle UV technology). Cells were then harvested at 4 °C, and the formation of crosslinked PhoQ/MgrB complex was then detected by the separation of total protein with polyacrylamide gel electrophoresis and Western blot with anti-Flag antibody. The crosslinked complex was purified by affinity chromatography, separated in an SDS polyacrylamide gel, and verified by mass spectrometry (see SI Materials and Methods for details).

### Molecular Modeling and Molecular Dynamics simulations

The PhoQ/MgrB complex was modeled using Alphafold2 multimer on Colab [PMID: 35637307]. We considered six different complexes: i. PhoQ, ii. PhoQ / MgrB (predictions ranked 1 to 3), iii. PhoQ / MgrB W20Y variant, and iv. PhoQ / MgrB W20A variant. Each complex was inserted into a bacterial membrane mimetic composed of a mixture of phosphatidylethanolamine (POPE) and phosphatidylglycerol (POPG) with a 3:1 ratio. The systems were solvated with a 17 Å water padding and neutralized with NaCl at a concentration of 150 mM. Each system was assembled using the CHARMM-GUI webserver [46].

The simulation was performed with the CHARMM36m force field, including CMAP corrections for the protein [47]. The water molecules were described with the TIP3P water parameterization [48]. The simulations were carried out with OPENMM molecular engine [49] following the minimization and equilibration protocols provided by CHARMM-GUI. The cutoff for non-bonded interactions was set to 12 Å with a switching distance at 10 Å. The periodic electrostatic interactions were computed using particle-mesh Ewald (PME) summation with a grid spacing smaller than 1 Å. Constant temperature of 310 K was imposed by Langevin dynamics with a damping coefficient of 1.0 ps. Constant pressure of 1 atm was maintained with Monte Carlo barostat [50]. The hydrogen mass repartitioning scheme was used to achieve a 4 fs time-step [51]. Each simulation was carried out up to 1 μs and up to 500 ns for the second and third ranked predictions of PhoQ / MgrB.

### Acceptor photobleaching FRET analysis

FRET measurements were performed as described before [30]. The expression of PhoQ-mNeonGreen and mCherry-MgrB was induced in *E. coli* lptD4213 strain with 0.008% arabinose and 10 μM IPTG, respectively. The cultures were grown to the mid-log phase (OD=0.6) in LB media. Cells were harvested and washed twice with pre-chilled tethering buffer (20 mM potassium phosphate pH 7.0, 1 μM methionine, and 10 mM lactic acid). The surface of a glass-bottom 96-well plate (Greiner) was treated with 0.1% poly-L-lysine for 15 minutes at room temperature, followed by removing poly-L-Lysine and air-dry for 5 minutes. The wells were washed twice using a pre-chilled tethering buffer. Subsequently, cells were incubated in the wells at room temperature for 25 minutes to allow attachment. Unattached cells were removed by washing with tethering buffer twice, and attached cells were overlaid with tethering buffer with or without 10 µg/ml C18G and incubated at room temperature for one hour. Acceptor photobleaching FRET was performed using the same microscope settings previously [30]. The details of the data acquisition and analysis can be found in the SI Materials and Methods.

## Acknowledgments

We thank L. Cassidy and A. Throley (the Christian-Albrecht University of Kiel) for performing mass spectrometry analyses, writing and editing the manuscript; H. Koch (the Albert Ludwig University of Freiburg) for pEVOL-pBpF vector and the site-directed photo-crosslinking protocol; M. Stopp for the disulfide crosslinking protocol; S. G. Sierra for technical assistance; members of the Yuan and the Sourjik laboratories for helpful discussions.

We thank G. Storz (National Institutes of Health), V. Sourjik (Max Planck Institute for Terrestrial Microbiology), R. Lill (Philipps University of Marburg), and S. Yadavalli (Rutgers University) for their suggestions to improve the manuscript.

This work is supported by the German Research Foundation (DFG) priority program 2002 YU 247/3-1 and the Max Planck Society (to J.Y.), the Swiss National Science Foundation (SNSF) grant PCEFP3_194606 (to T.L. and S.L.).

J.Y. and T.L. designed the experiments; S.J., L.C.S., C.K., S.L., and J.Y. performed the experiments; S.J., G.M., L.C.S., C.K., S.L., T.L., V.S., and J.Y. analyzed data; J.Y., T.L., and S.J. wrote the manuscript.

## Supporting Information

### SI Materials and Methods

#### Cloning and plasmid construction

*E. coli phoQ-his_6_* gene was cloned into pET-Duet-1 vector at NcoI/BamHI restriction sites, and *E. coli flag-mgrB* gene was cloned into pET-Duet-1 vector at NdeI/ EcoRV restriction sites. The pEVOL-pBpF vector is a gift from Dr. Hans-Georg Koch (Freiburg, Germany). Mutations in *phoQ* and *mgrB* were generated using the Q5 site-directed mutagenesis kit (New England BioLabs) following the manufacturer’s instruction. All constructs were verified by DNA sequencing.

#### Purification of the crosslinked PhoQ-His/FLAG-MgrB complex

A two-liter culture was prepared for the purification of the crosslinked PhoQ-His/FLAG-MgrB complex. After UV crosslinking, cells were harvested, resuspended in the resuspension buffer (20 mM Tris-HCl, pH 7.5, 300 mM NaCl, 0.1 mM PMSF), then lysed with LM10 microfluidizer at 4°C. The lysate was centrifuged at 11,000 for 10 min to remove cell debris. The supernatant was collected and centrifuged at 13,500 g for 2 hours to pellet the total membrane, which was then dissolved in the resuspension buffer containing 1% (wt/vol) n-dodecyl-β-D-maltopyranoside (DDM) with gentle shaking at 4 °C overnight. Insoluble fraction was spun down by centrifugation at 21,000 g for 45 min. The supernatant was loaded onto a TALON column (Cytiva), which was then washed three times with buffer containing 50 mM Tris-HCl, pH 8.0, 500 mM NaCl, 10% glycerol, 10 mM imidazole, 0.1 mM PMSF and 0.03% DDM. The protein was eluted with elution buffer containing 50 mM Tris-HCl, pH 8.0, 500 mM NaCl, 10% glycerol, 200 mM imidazole, 0.1 mM PMSF and 0.03% DDM, and concentrated to ∼200 µL. The elution was then incubated with anti-FLAG M2 magnetic beads (Sigma) at 4 °C overnight. The magnetic beads were washed with Tris buffered saline, and then incubated with 3 X FLAG peptide solution (Sigma) at room temperature for one hour to elute FLAG-tagged proteins. The supernatant was collected, concentrated to ∼50 µl, and analyzed using gel electrophoresis with a 7.5% tris-glycine polyacrylamide gel.

#### Protein verification with mass spectrometry

Crosslinked and control bands were excised from SDS-PAGE gels and de-stained and sliced into approx. 1mm^3^ pieces. Proteins in the gel pieces were reduced (10 mM dithiothreitol, 56°C for 30 min), and alkylated (50 mM chloroacetamide, 30 min, RT, in the dark), prior to overnight enzymatic hydrolysis with chymotrypsin (Promega, WI, USA) (0.1 µg of chymotrypsin per band in 50 mM ammonium bicarbonate buffer pH 7.4, 2 mM CaCl2) at 37°C. Following overnight digestion, the peptides were collected and dried down via vacuum centrifugation (Concentrator Plus, Eppendorf). Peptides were resuspended in 10 µl of HPLC loading buffer (3 % ACN + 0.1 % trifluoroacetic acid) prior to LC-MS/MS analysis.

Chromatographic separation was performed on a Dionex U3000 nanoHPLC system (Thermo, Germany) equipped with an Acclaim pepmap100 C18 column (2 μm particle size, 75 μm × 500 mm) coupled online to a QExactive Plus mass spectrometer (Thermo, Bremen). The separation was performed over a 60-minute run: 5 % B for 3 minutes followed by linear gradients from 5 % to 50 % B over 30 minutes, then 50 % to 90 % over 1 minute, and 10 minutes at 90 % B. Inter-run equilibration was achieved by 15 minutes at 5 % B. Eluent A: 0.05 % formic acid (FA), eluent B: 80 % ACN + 0.04 % FA. A flow rate of 300 nl/min was used and 1 μl of sample was injected per run. For crosslink samples a second injection of 6 µl was also performed. Full scan MS acquisition was performed (300-1500 m/z, resolution 70,000, AGC target 3e6, max injection time (IT) 100 ms) with subsequent data dependent MS/MS of the top 10 most intense ions via HCD ion activation at NCE 27 (resolution 17,500, AGC target 1e5, isolation window 1.6 m/z, max IT 50 ms); dynamic exclusion (20 s duration) and lock mass at 445.12003 m/z were enabled.

For protein identifications, the MS data files were processed with the Proteome Discoverer^TM^ software suite (Ver. 2.5.0.400) (Thermo, Germany). Raw files were searched using the SequestHT algorithmand Target Decoy PSM validator node, against a combined database that included the proteins of interest (PhoQ-Histagged, the modified MgrB-pBpA incorporation variants), the UniProt canonical proteins for *Escherichia coli* (strain K12) (accessed: 2022.03.10), and common laboratory contaminants (cRAP list). Searches were performed with full chymotrypsin specificity, and a maximum 3 missed cleavages. Precursor mass tolerance 10 ppm. Fragment mass tolerance 0.02 Da. Fixed modification of carbamidomethyl (Cys), dynamic modification oxidation (Met). Strict parsimony criteria were applied: Target FDR <1% was applied, and at least one unique high confidence peptide was required for identification. For the small protein MgrB, manual assessment of the peptide spectral matches was also performed.

#### Acceptor photobleaching FRET data acquisition and analysis

For each well, we acquired three or more sequences of images on isolated fields of view with the following protocol: (a) 2 images in the acceptor channel; (b) 140 images in the donor channel followed by (c) 16 seconds acceptor photobleaching (no image acquisition); (d) 50 images in the donor channel; and then (e) 2 images in the acceptor channel. FRET efficiency was calculated as the increase of donor fluorescence signal upon acceptor photobleaching divided by the total donor signal after acceptor photobleaching. In order to correct for the donor photobleaching present during steps b to d, we performed linear fitting (RStudio) of the donor fluorescence signal versus time for both pre- and postbleaching curves.

### SI Figures and Tables

**Fig. S1.**
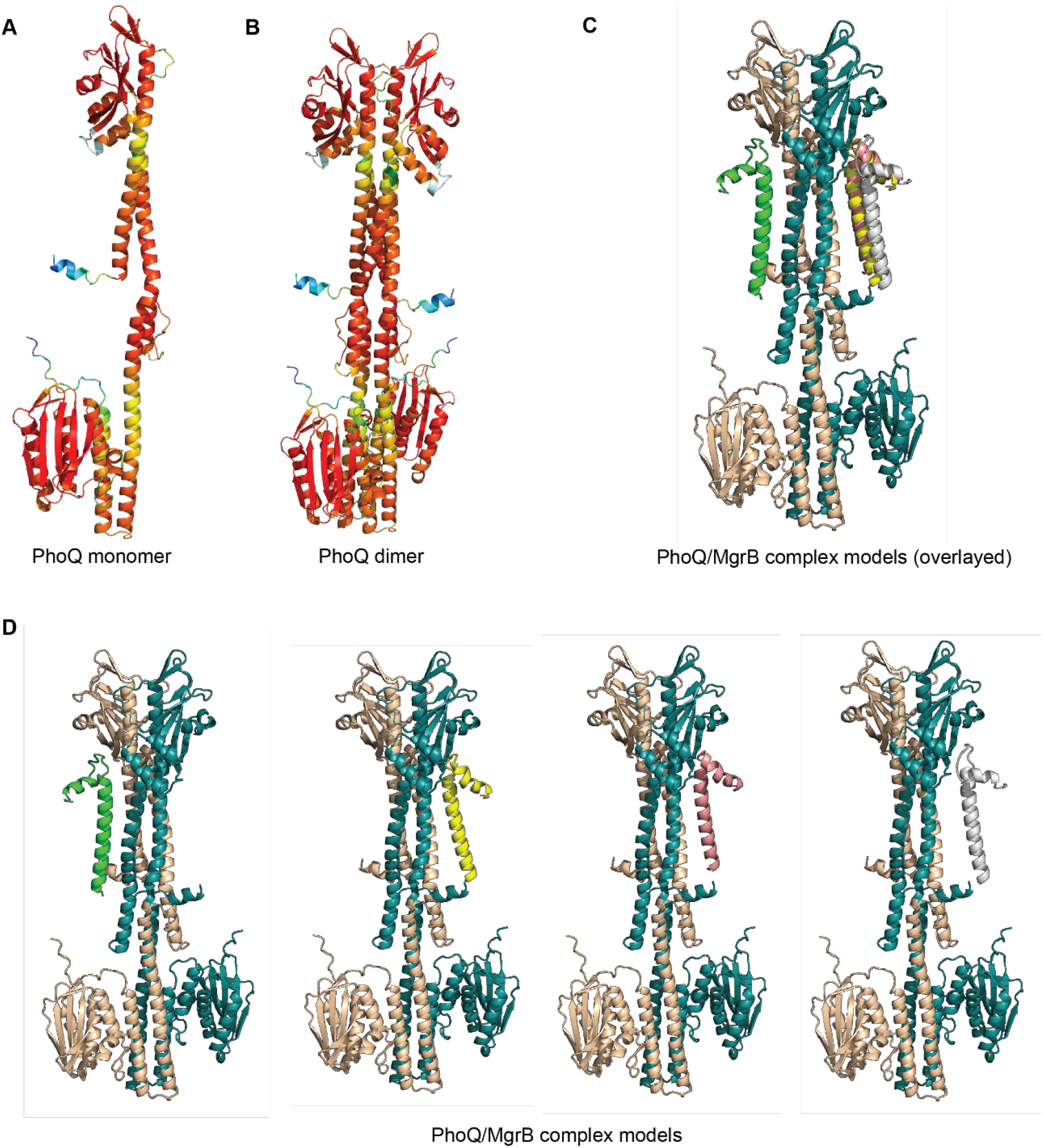
Structural models predicted by Alphafold2. (**A-B**) The predicted PhoQ monomer and dimer are colored according to the pLDDT score to show the regions with lower prediction confidence. Red indicates highest confidence and blue indicates lowest confidence. The four predicted PhoQ/MgrB complex models ranked from two to five by Alphafold2. Models are shown with a PhoQ dimer (colored in wheat and deep teal) and four MgrB molecules overlayed (**C**) or separated (**D**) in green, yellow, pink and grey ranked from 2 to 5, respectively. All structural figures were prepared using PyMOL unless otherwise stated.

**Fig. S2.**
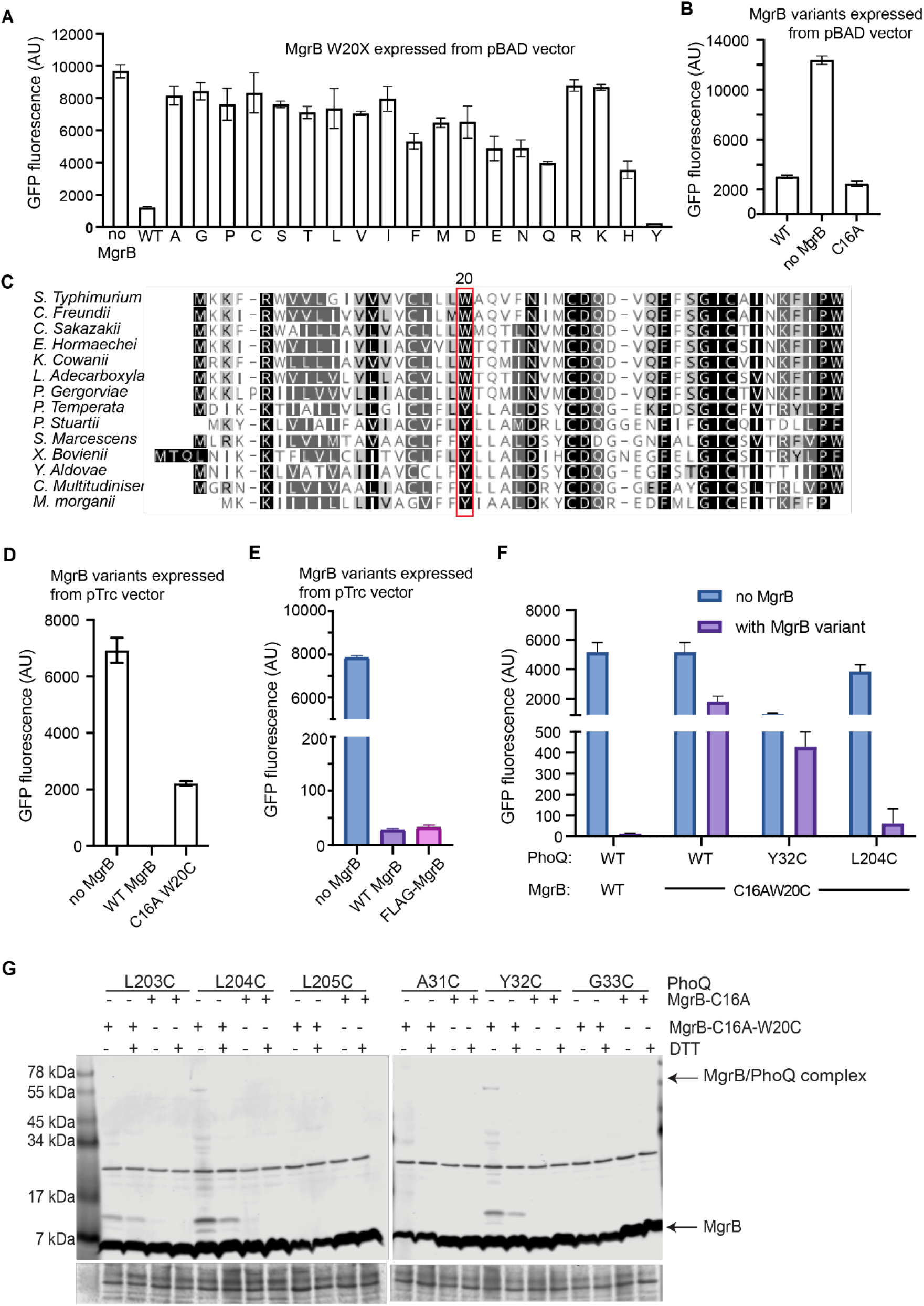
The function of MgrB variants and the crosslinking with PhoQ from MgrB position 20. The activity analysis of MgrB W20 variants (**A**) and MgrB C16A (**B**) using the reporter plasmid pUA66 *P_mgtLA_*-*gfp*. The *E. coli ΔmgrB* strain harboring the reporter plasmid and the pBAD plasmid with *mgrB* variants were grown to the early log phase in LB media supplemented with 0.008% arabinose. GFP fluorescence in cells was then measured by flow cytometry. (**C**) Sequence alignment of MgrB homologs from fourteen bacterial species. The conservation of the amino acid at position 20 (*E. coli* numbering) is highlighted with a red rectangle. (**D**) The functional assay of MgrB variants expressed from pTrc99A vector. The pUA66 P*_mgtLA_*-*gfp*, pBAD33 *phoQ*, and pTrc99A plasmid with flag tagged *mgrB* variants, were transformed into *E. coli ΔphoQΔmgrB* cells. The resulting transformants were grown overnight at 37 °C in LB medium supplemented with 10 mM MgSO4, then diluted 1:100 to fresh LB medium supplemented with 1 mM MgSO4, 0.008% arabinose, 10 µM IPTG, and antibiotics when appropriate. The cultures were grown at 37 °C with vigorous shaking till the early log phase. The GFP fluorescence of cells was then monitored by flow cytometry. (**E**) The functional assay of MgrB and Flag-MgrB expressed from pTrc99A vector. The pUA66 P*_mgtLA_*-*gfp* and pTrc99A plasmid with *mgrB* or *flag-mgrB*, were transformed into *E. coli ΔmgrB* cells. The resulting transformants were grown overnight at 37 °C in LB medium supplemented with 10 mM MgSO4, then diluted 1:100 to fresh LB medium supplemented with 1 mM MgSO4, 10 µM IPTG, and antibiotics when appropriate. The cultures were grown at 37 °C with vigorous shaking till the early log phase. The GFP fluorescence of cells was then monitored by flow cytometry. (**F**) The functional assay of the indicated PhoQ and MgrB variants using the same protocol as in D. (**G**) Western blot analysis of *E. coli* membrane extracts after disulfide crosslinking. The indicated PhoQ and FLAG-tagged MgrB variants were expressed in *E. coli*, followed by Cu-phenanthroline catalyzed disulfide crosslinking (details in Materials and Methods). The total protein stain of the PVDF membranes serves as loading controls. Data are representative of at least three independent experiments. In A, B, D, E and F, each data point is shown with the calculated average and standard deviation from three independent biological replicates.

**Fig. S3.**
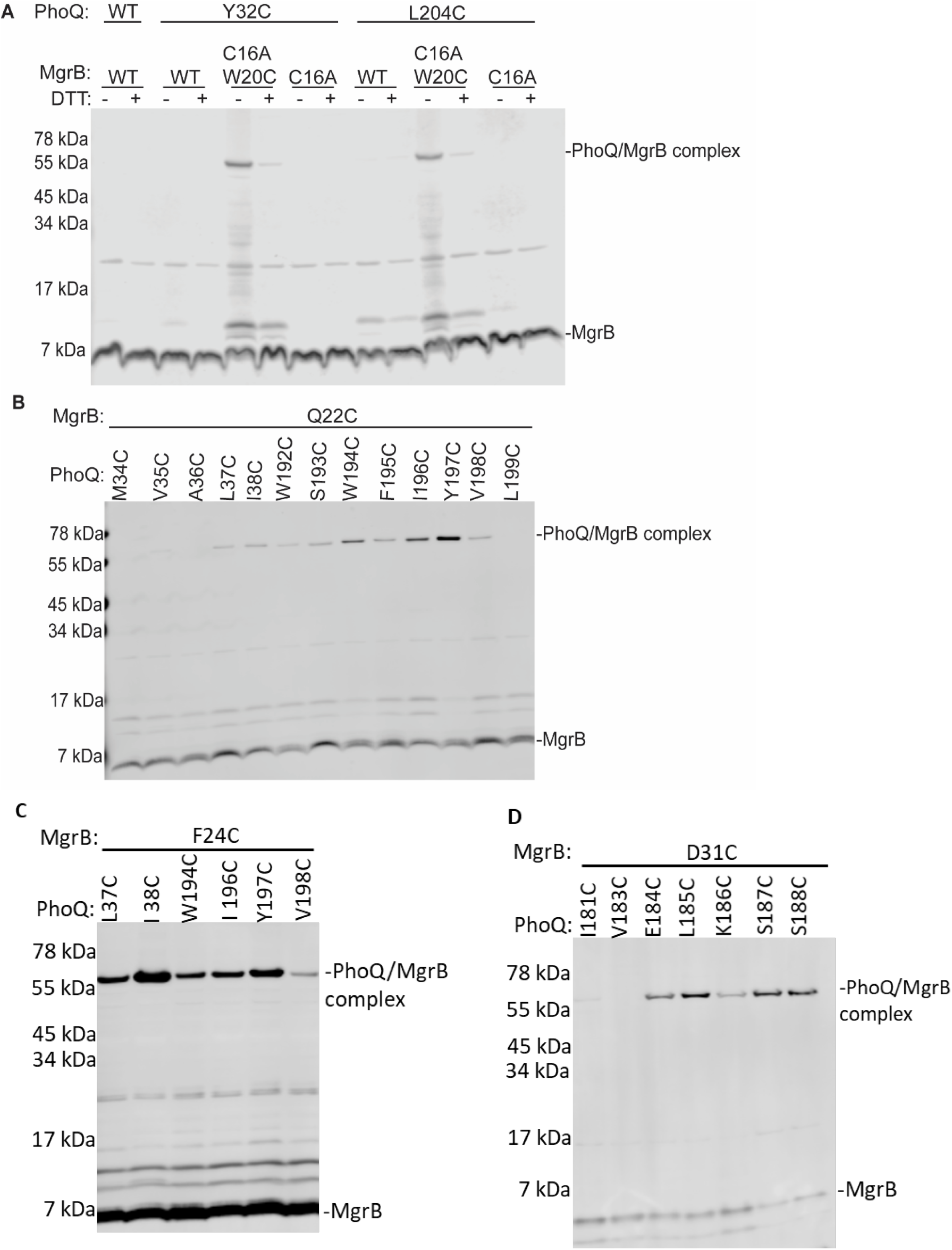
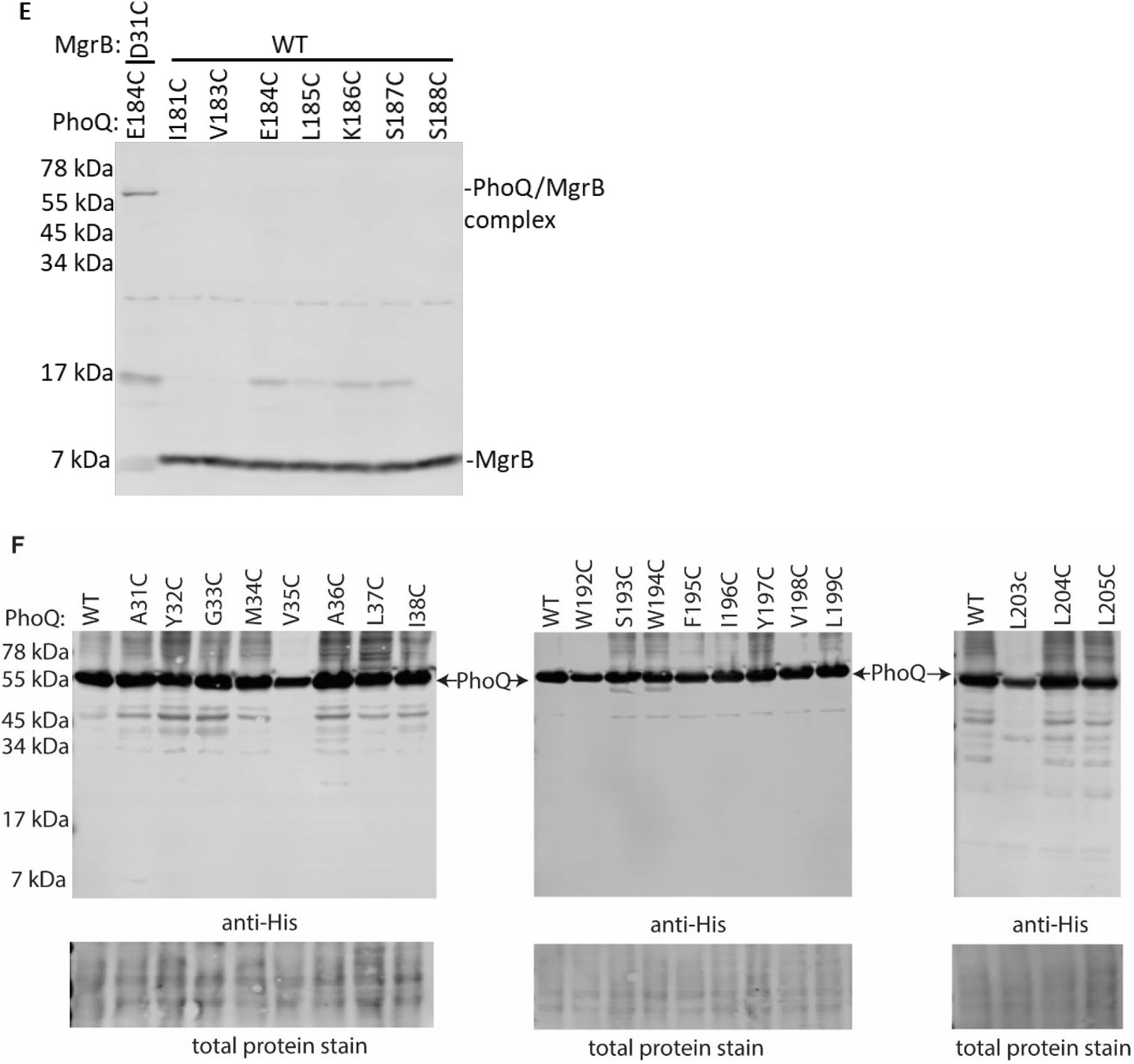
The full Western blot results of the figures in the main text. (**A**) is the full blot of Fig. 1D. (**B**) is the full blot of Fig. 1E. (**C**) is the full blot of Fig. 1F. (**D, E**) are the full blots of Fig. 2B. Data are representative of at least three independent experiments. (**F**) Western blot analysis of the expression level of PhoQ variants. The indicated PhoQ-His variants were expressed in *E. coli* by inducing with 0.008% arabinose. Cells were harvested in the mid log phase and the total membrane was extracted for the western blot analysis. The total protein stain of the PVDF membranes serves as a loading control. Data are representative of two independent experiments.

**Fig. S4.**
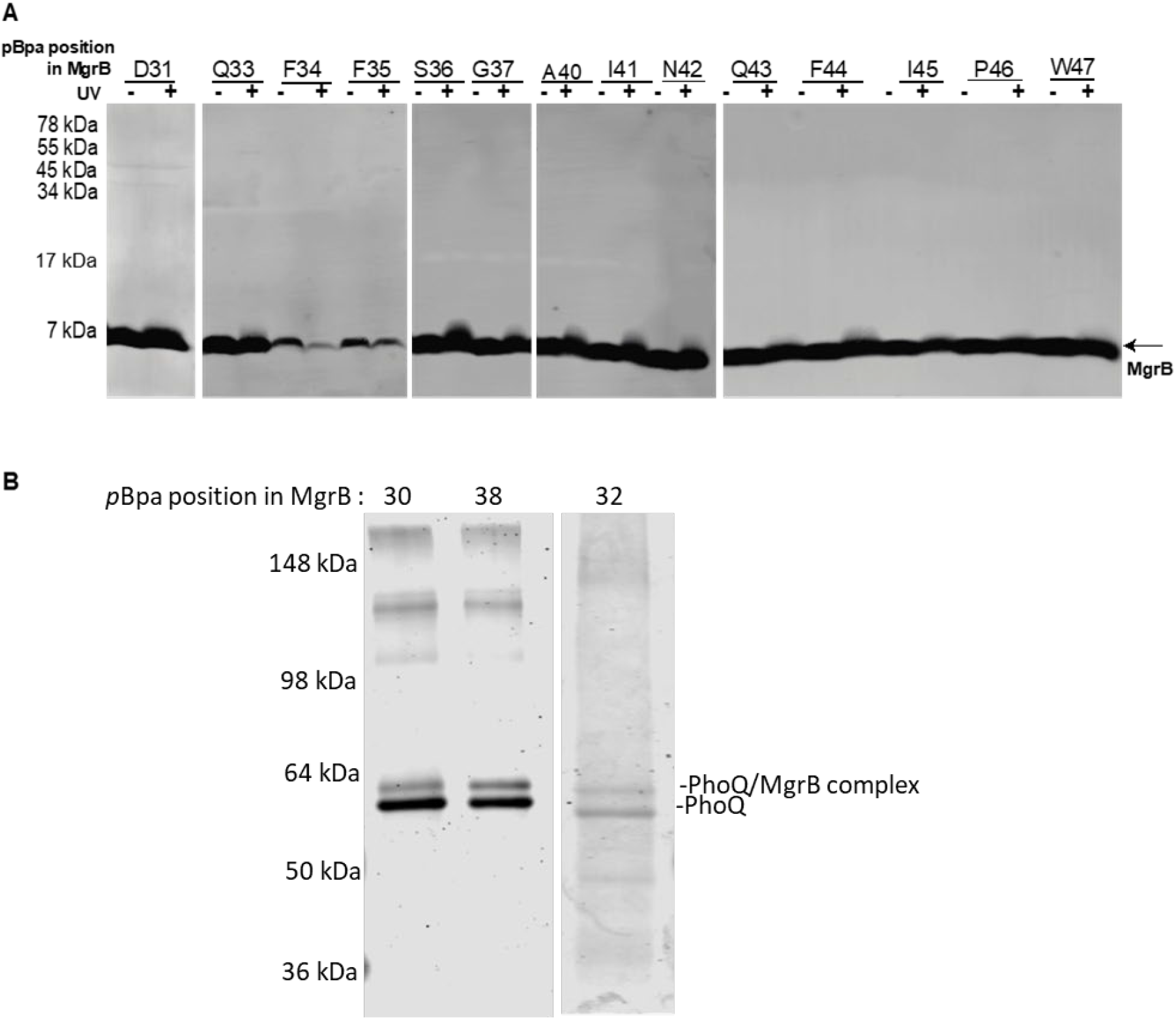
Mapping the binding interface of MgrB on PhoQ by using site-specific UV crosslinking in *vivo*. (**A**) *E. coli* BL21 DE3 cells expressing PhoQ-His_6_, FLAG-MgrB or *p*Bpa-containing FLAG-MgrB variants were irradiated with or without UV light. The cells were lysed in SDS sample buffer. Proteins were separated on 7.5% tris-glycine polyacrylamide gels and transfer to PVDF membranes. FLAG tagged proteins were detected with an anti-FLAG primary antibody and an IRDye 800CW-conjugated secondary antibody. Data are representative of three independent experiments. (**B**) Purification of PhoQ-His/FLAG-MgrB crosslinked complex. Elution from a two-step affinity purification was separated on 7.5% tris-glycine gels. The protein bands in the gel were visualized with the ready-blue protein stain. Data are representative of two independent experiments.

**Fig. S5.**
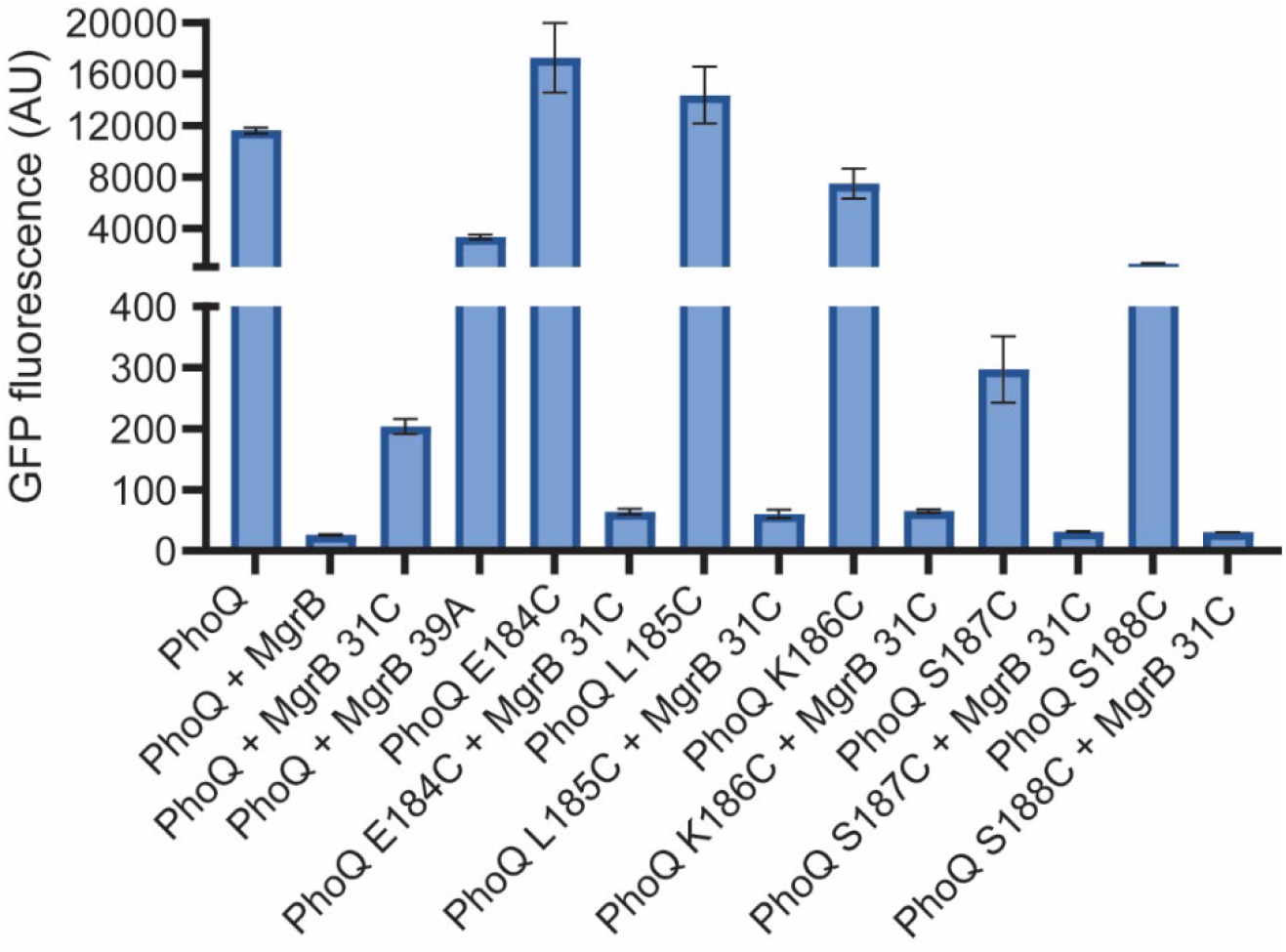
The functional assay of PhoQ variants with or without MgrB. *E. coli* MG1655 *ΔphoQΔmgrB* cells harboring pBAD33 *phoQ* variants, the GFP reporter plasmid pUA66 P*_mgtA_*-*gfp*, pTrc99A with or without *mgrB* variants were grown in LB supplemented with 0.008% arabinose and 10 µM IPTG till the early-log phase. The GFP fluorescence of the cells was measured with flow cytometry. Each data point is shown with the calculated average and standard deviation from three biological replicates. The results are the representative of at least three independent experiments.

**Fig. S6.**
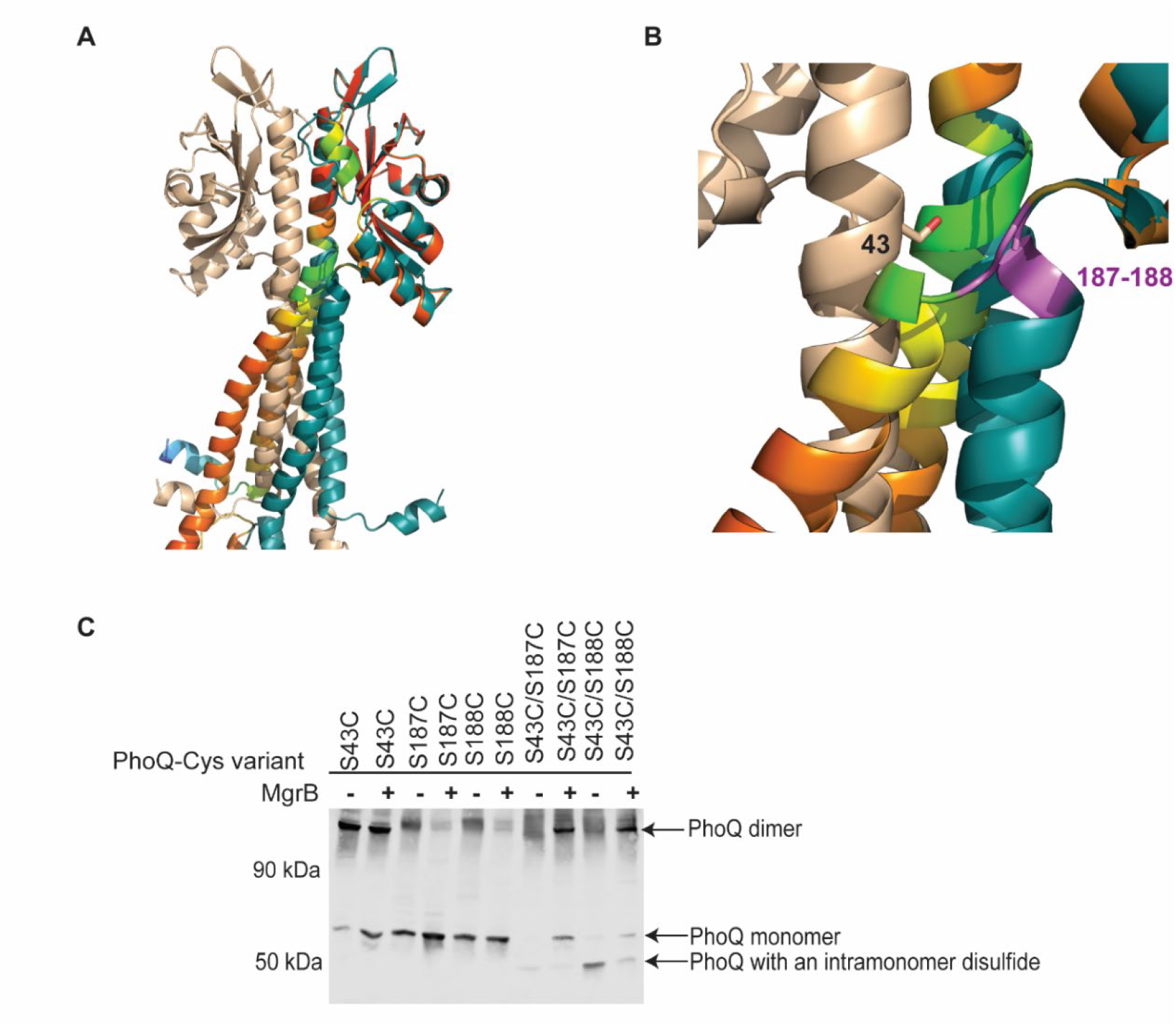
The conformational change of PhoQ linker region in the presence of MgrB. (**A**) The structural superimposition of the periplasmic and TM domains of the PhoQ dimer (wheat and deep teal) and monomer (pLDDT) is shown. The periplasmic domain was used for structural alignment in Pymol. (**B**) Zoom in view of the linker region of PhoQ. Residue 43 is shown in ball- and-stick. Residues 187 and 188 are colored in purple in both monomer and dimer structures. (**C**) Western blot analysis of disulfide crosslinking within the PhoQ molecule in the presence or absence of MgrB. The crosslinked PhoQ species was detected with an anti-His antibody. Data are representative of at least three independent experiments.

**Fig. S7.**
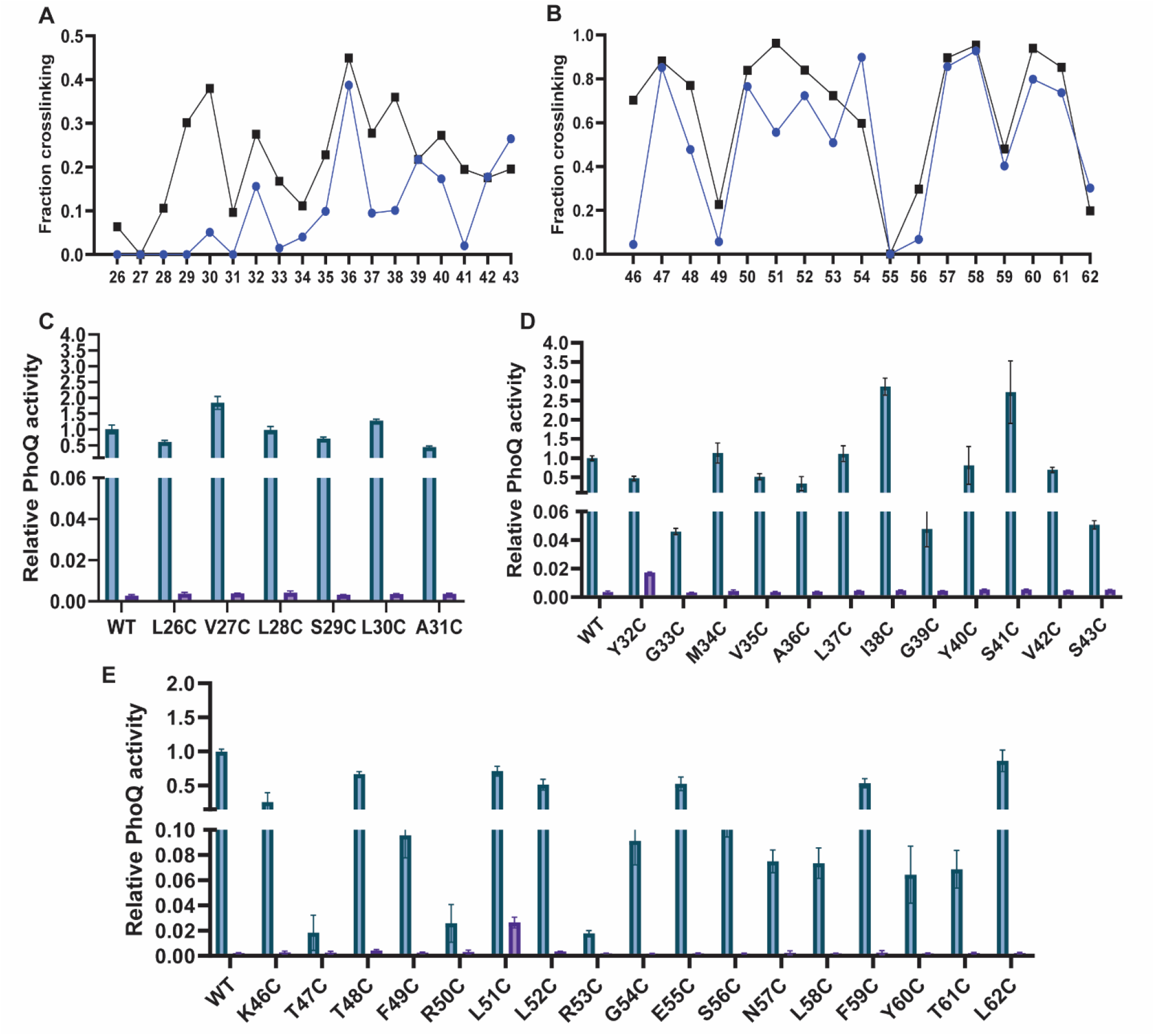
PhoQ_Cys variants in the presence or absence of MgrB. (**A-B**) Quantification of the disulfide crosslinking efficiency of PhoQ_Cys variants in the presence (blue curve) and absence (black curve) of MgrB. The PhoQ dimer and monomer bands in Fig. 4C were quantified with ImageJ. The fraction of crosslinked dimer was calculated using the formula: crosslinking fraction = dimer / (dimer + monomer). (**C-E**) Functional assays of PhoQ_Cys variants with (blue) and without (purple) MgrB. *E. coli ΔphoQΔmgrB* cells harboring pUA66 P*_mgtA_*-*gfp*, pBAD33 *phoQ* variants, and pTrc99A empty or pTrc99A *mgrB* were grown overnight at 37 °C in LB medium supplemented with 10 mM MgSO_4_. Cultures were then diluted 1:100 to fresh LB medium supplemented with 1 mM MgSO_4_, 0.008% arabinose and 10 µM IPTG, and grown at 37 °C with vigorous shaking till the early log phase (OD=0.4-0.5). The fluorescence of cells was monitored with flow cytometry and normalized to the PhoQ wild type. Each data point in C, D and E is shown with the calculated average and standard deviation from three biological replicates. The results are the representative of three independent experiments.

**Fig. S8.**
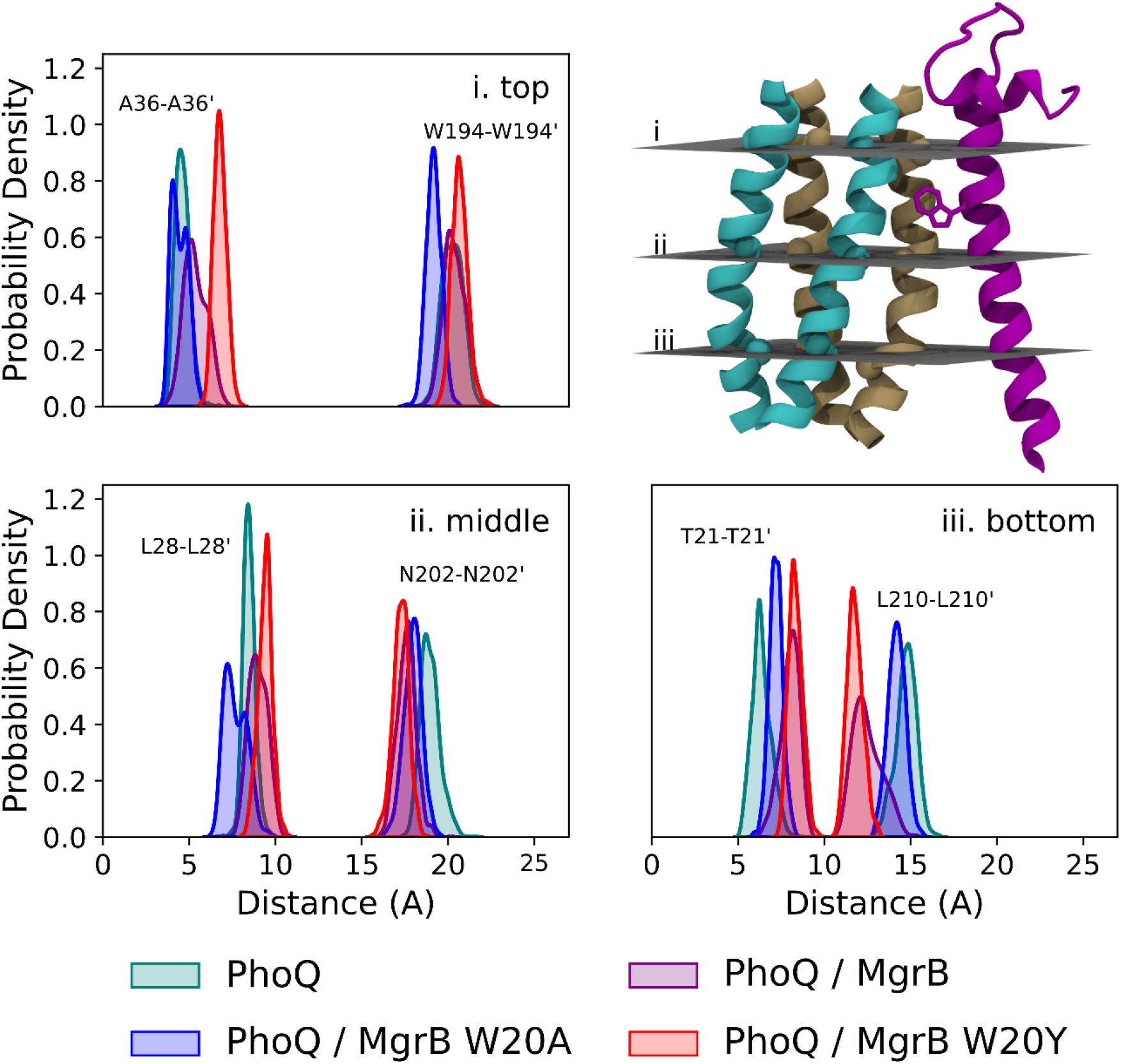
MD simulations of PhoQ/MgrB complex models. Distribution in distances between TM1 – TM1’ (left) and TM2 – TM2’ (right) at the top, middle, and bottom of PhoQ TM domain.

**Fig. S9.**
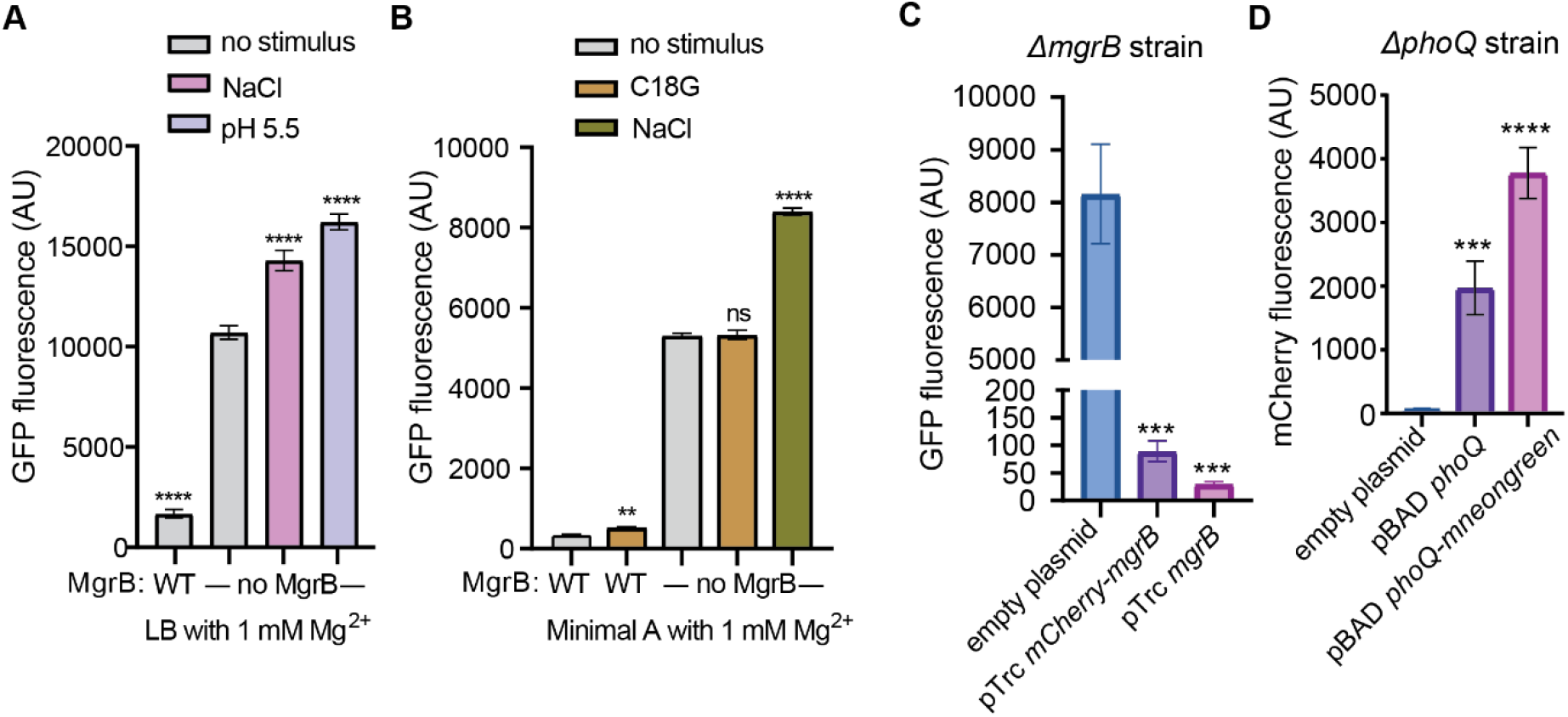
The functional assays with fluorescent reporters. PhoQ activity was monitored with a GFP reporter plasmid (pUA66 P*_mgtLA_-gfp*) with or without MgrB in LB (**A**) and in minimal A medium (**B**) supplemented with 1 mM magnesium. Breifly, *E. coli ΔmgrB* strain harboring the GFP reporter plasmid and a pBAD plasmid with or without the wild-type mgrB gene, was grown in the presence of the indicated PhoQ stimuli to early log phase. The GFP fluorescence of cells was then measured by flow cytometry. Data are representative of at least three independent experiments. The function of fusion proteins mCherry-MgrB (**C**) and PhoQ-mneonGreen (**D**) were anlalyzed with the GFP report and a mCherry reporter (pUA66 P*_mgtLA_-mcherry*). Indicated deletion strains were complemented with a plasmid harboring the gene of indicated fusion proteins. The expression of the fusion protein was induced with 10 µM IPTG and 0.008% arabinose for the pTrc and pBAD plasmids, respectively. Cells were grown to early log phase and the fluorescence of cells was measured by flow cytometry. Cells transformed with empty plasmids and plasmids harboring the corresponding wild-type genes were used as negative and positive controls, respectively. Data are representative of two independent experiments.

**Table S1.** List of strains used in this study.

| Strain | Genotype | Source or Reference |
| --- | --- | --- |
| JingY11 | MG1655 | *E. coli* Genetic Stock Center, CGSC# 7740 |
| lptD4213 | F<sup>-</sup>, *lptD4213*, *galK2*(Oc), $\lambda^-$, *IN(rrnD-rrnE)1*, *rpsL200*(strR), *rph-1* | Yale Coli Genetic Stock Center |
| *E. coli* BL21(DE3) | BL21(DE3) | New England Biolabs |
| JingY34 | MG1655 $\Delta mgrB::FRT$ | [1] |
| JingY90 | MG1655 $\Delta phoQ::FRT \Delta mgrB::FRT$ | [1] |

**Table S2.** List of plasmids used in this study.

| Plasmid | Genotype | Source or Reference |
| --- | --- | --- |
| pJY1 | pBAD33 *phoQ*-(*his*)<sub>6</sub> | This study |
| pJY31 | pBAD33RBS | [2] |
| pJY151 | pBAD33RBS *mgrB* | [1] |
| pOB2 | pTrc99A *mcherry* | [1] |
| pJY569 | pTrc99A *mcherry-mgrB* | [1] |
| pJY573 | pBAD33RBS *phoQ-mneongreen* | [1] |
| pJY317 | pTrc99A *flag-mgrB* | This study |
| pJS1 | pET-Duet-1 EC *phoQ-his*<sub>6</sub>, *flag-mgrB* | This study |
| pEVOL | pEVOL-pBpF | A gift from Dr. Hans-Georg Koch |
| pLS1 | pBAD33RBS *phoQ* L26C | This study |
| pLS2 | pBAD33RBS *phoQ* V27C | This study |
| pLS3 | pBAD33RBS *phoQ* L28C | This study |
| pLS4 | pBAD33RBS *phoQ* S29C | This study |
| pLS5 | pBAD33RBS *phoQ* L30C | This study |
| pLS6 | pBAD33RBS *phoQ* A31C | This study |
| pLS7 | pBAD33RBS *phoQ* Y32C | This study |
| pLS8 | pBAD33RBS *phoQ* G33C | This study |
| pLS9 | pBAD33RBS *phoQ* M34C | This study |
| pLS10 | pBAD33RBS *phoQ* V35C | This study |
| pLS11 | pBAD33RBS *phoQ* A36C | This study |
| pLS12 | pBAD33RBS *phoQ* L37C | This study |
| pLS13 | pBAD33RBS *phoQ* I38C | This study |
| pLS14 | pBAD33RBS *phoQ* G39C | This study |
| pLS15 | pBAD33RBS *phoQ* Y40C | This study |
| pLS16 | pBAD33RBS *phoQ* S41C | This study |
| pLS17 | pBAD33RBS *phoQ* V42C | This study |
| pLS18 | pBAD33RBS *phoQ* S43C | This study |
| pLS19 | pBAD33RBS *phoQ* F44C | This study |
| pJS4 | pBAD33RBS *phoQ* K46C | This study |
| pJS5 | pBAD33RBS *phoQ* T47C | This study |
| pJS6 | pBAD33RBS *phoQ* T48C | This study |
| pJS7 | pBAD33RBS *phoQ* F49C | This study |
| pJS8 | pBAD33RBS *phoQ* R50C | This study |
| pJS9 | pBAD33RBS *phoQ* L51C | This study |
| pJS10 | pBAD33RBS *phoQ* L52C | This study |
| pJS11 | pBAD33RBS *phoQ* R53C | This study |
| pJS12 | pBAD33RBS *phoQ* G54C | This study |
| pJS13 | pBAD33RBS *phoQ* E55C | This study |
| pJS14 | pBAD33RBS *phoQ* S56C | This study |
| pJS15 | pBAD33RBS *phoQ* N57C | This study |
| pJS16 | pBAD33RBS *phoQ* L58C | This study |
| pJS17 | pBAD33RBS *phoQ* F59C | This study |
| pJS18 | pBAD33RBS *phoQ* Y60C | This study |
| pJS19 | pBAD33RBS *phoQ* T61C | This study |
| pJS20 | pBAD33RBS *phoQ* L62C | This study |
| pJS21 | pBAD33RBS *phoQ* D179C | This study |
| pJS22 | pBAD33RBS *phoQ* T181C | This study |
| pJS23 | pBAD33RBS *phoQ* I183C | This study |
| pJS24 | pBAD33RBS *phoQ* E184C | This study |
| pJS25 | pBAD33RBS *phoQ* L185C | This study |
| pJS26 | pBAD33RBS *phoQ* K186C | This study |
| pLS20 | pBAD33RBS *phoQ* S187C | This study |
| pLS21 | pBAD33RBS *phoQ* S188C | This study |
| pLS22 | pBAD33RBS *phoQ* S43C-S187C | This study |
| pLS23 | pBAD33RBS *phoQ* S43C-S188C | This study |
| pCK1 | pBAD33RBS *phoQ* S193C | This study |
| pCK2 | pBAD33RBS *phoQ* W194C | This study |
| pCK3 | pBAD33RBS *phoQ* F195C | This study |
| pCK4 | pBAD33RBS *phoQ* I196C | This study |
| pCK5 | pBAD33RBS *phoQ* Y197C | This study |
| pCK6 | pBAD33RBS *phoQ* V198C | This study |
| pCK7 | pBAD33RBS *phoQ* L199C | This study |
| pCK8 | pBAD33RBS *phoQ* S200C | This study |
| pCK9 | pBAD33RBS *phoQ* N202C | This study |
| pCK10 | pBAD33RBS *phoQ* L203C | This study |
| pCK11 | pBAD33RBS *phoQ* L204C | This study |
| pCK12 | pBAD33RBS *phoQ* L205C | This study |
| pCK13 | pBAD33RBS *phoQ* V206C | This study |
| pCK14 | pBAD33RBS *phoQ* I207C | This study |
| pCK15 | pBAD33RBS *phoQ* L209C | This study |
| pCK16 | pBAD33RBS *phoQ* L210C | This study |
| pCK17 | pBAD33RBS *phoQ* W211C | This study |
| pCK18 | pBAD33RBS *phoQ* V212C | This study |
| pCK19 | pBAD33RBS *phoQ* A213C | This study |
| pCK20 | pTrc99A *mgrB* C16A | This study |
| pCK21 | pTrc99A *mgrB* C16A W20C | This study |
| pCK22 | pTrc99A *mgrB* C16A Q22C | This study |
| pCK23 | pTrc99A *mgrB* C16A F24C | This study |
| pJS27 | pTrc99A *mgrB* D31C | This study |
| pJY268 | pBAD33RBS *mgrB* W20A | This study |
| pJY156 | pBAD33RBS *mgrB* C28A | This study |
| pJS51 | pET-Duet-1 *mgrB* N25-TAG | This study |
| pJS52 | pET-Duet-1 *mgrB* M26-TAG | This study |
| pJS53 | pET-Duet-1 *mgrB* M27-TAG | This study |
| pJS54 | pET-Duet-1 *mgrB* D29-TAG | This study |
| pJS55 | pET-Duet-1 *mgrB* Q30-TAG | This study |
| pJS56 | pET-Duet-1 *mgrB* D31-TAG | This study |
| pJS57 | pET-Duet-1 *mgrB* V32-TAG | This study |
| pJS58 | pET-Duet-1 *mgrB* Q33-TAG | This study |
| pJS59 | pET-Duet-1 *mgrB* F34-TAG | This study |
| pJS60 | pET-Duet-1 *mgrB* F35-TAG | This study |
| pJS61 | pET-Duet-1 *mgrB* S36-TAG | This study |
| pJS62 | pET-Duet-1 *mgrB* G37-TAG | This study |
| pJS63 | pET-Duet-1 *mgrB* I38-TAG | This study |
| pJS64 | pET-Duet-1 *mgrB* A40-TAG | This study |
| pJS65 | pET-Duet-1 *mgrB* I41-TAG | This study |
| pJS66 | pET-Duet-1 *mgrB* N42-TAG | This study |
| pJS67 | pET-Duet-1 *mgrB* Q43-TAG | This study |
| pJS68 | pET-Duet-1 *mgrB* F44-TAG | This study |
| pJS69 | pET-Duet-1 *mgrB* I45-TAG | This study |
| pJS70 | pET-Duet-1 *mgrB* P46-TAG | This study |
| pJS71 | pET-Duet-1 *mgrB* W47-TAG | This study |

**Table S3.**
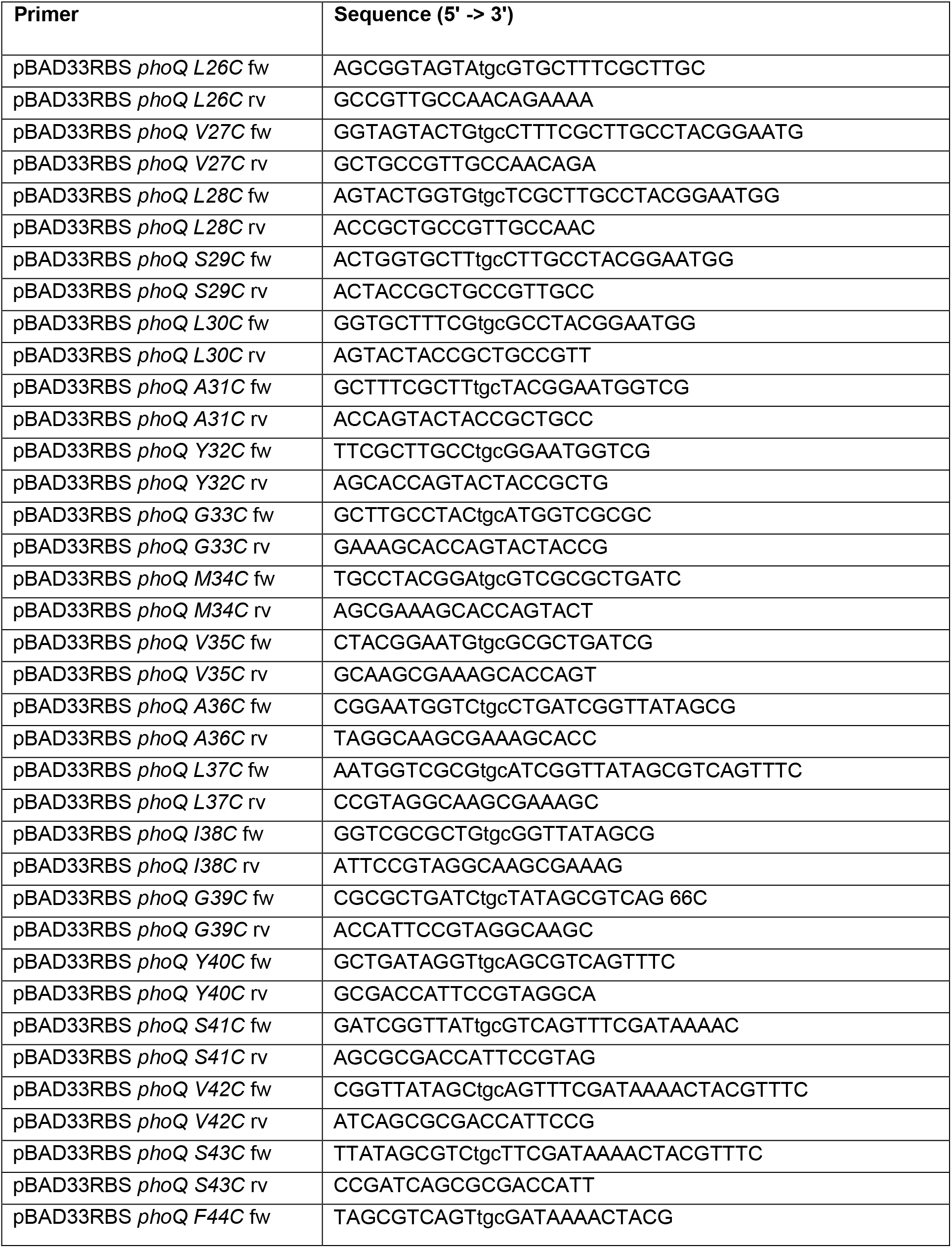
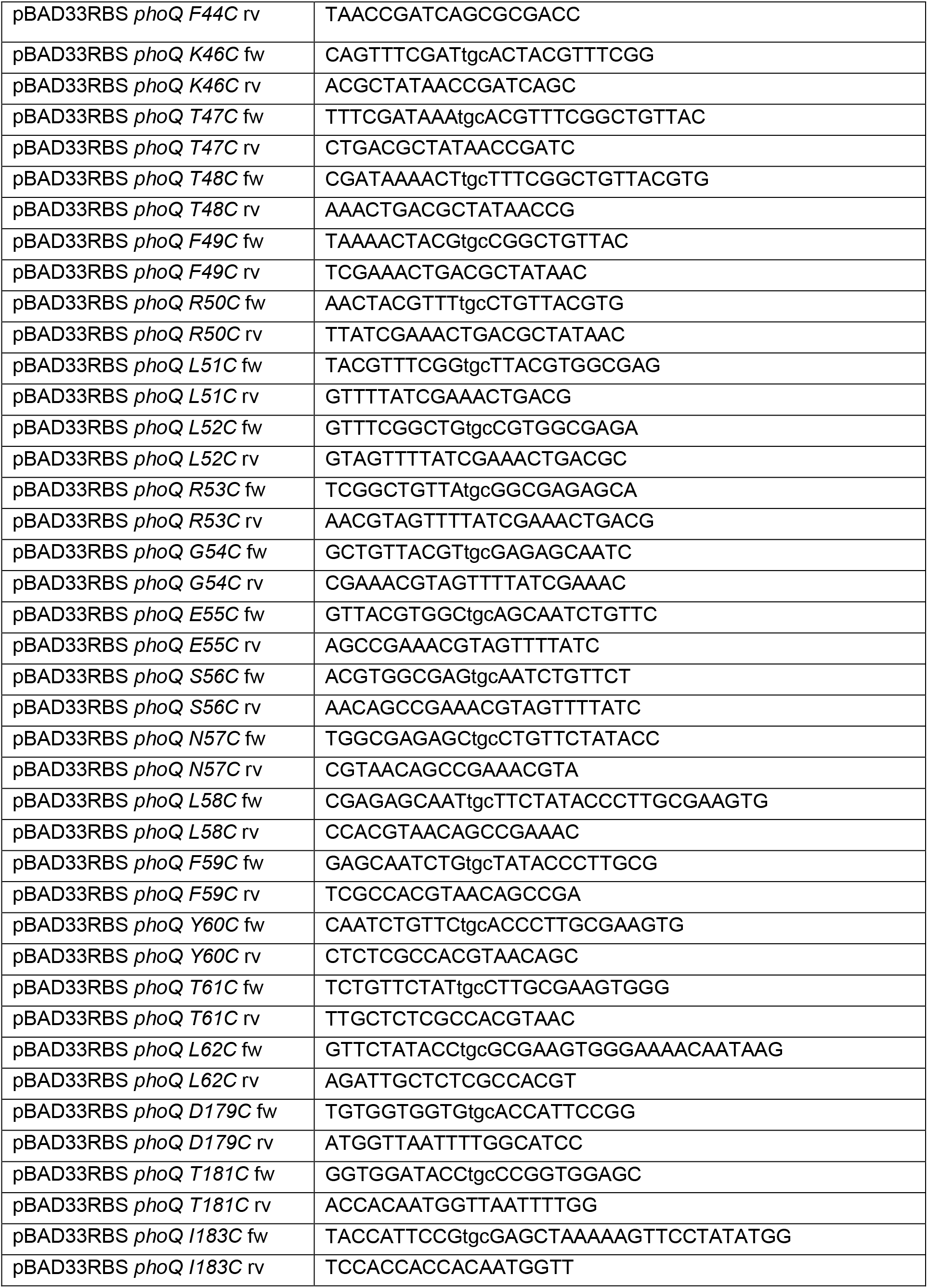
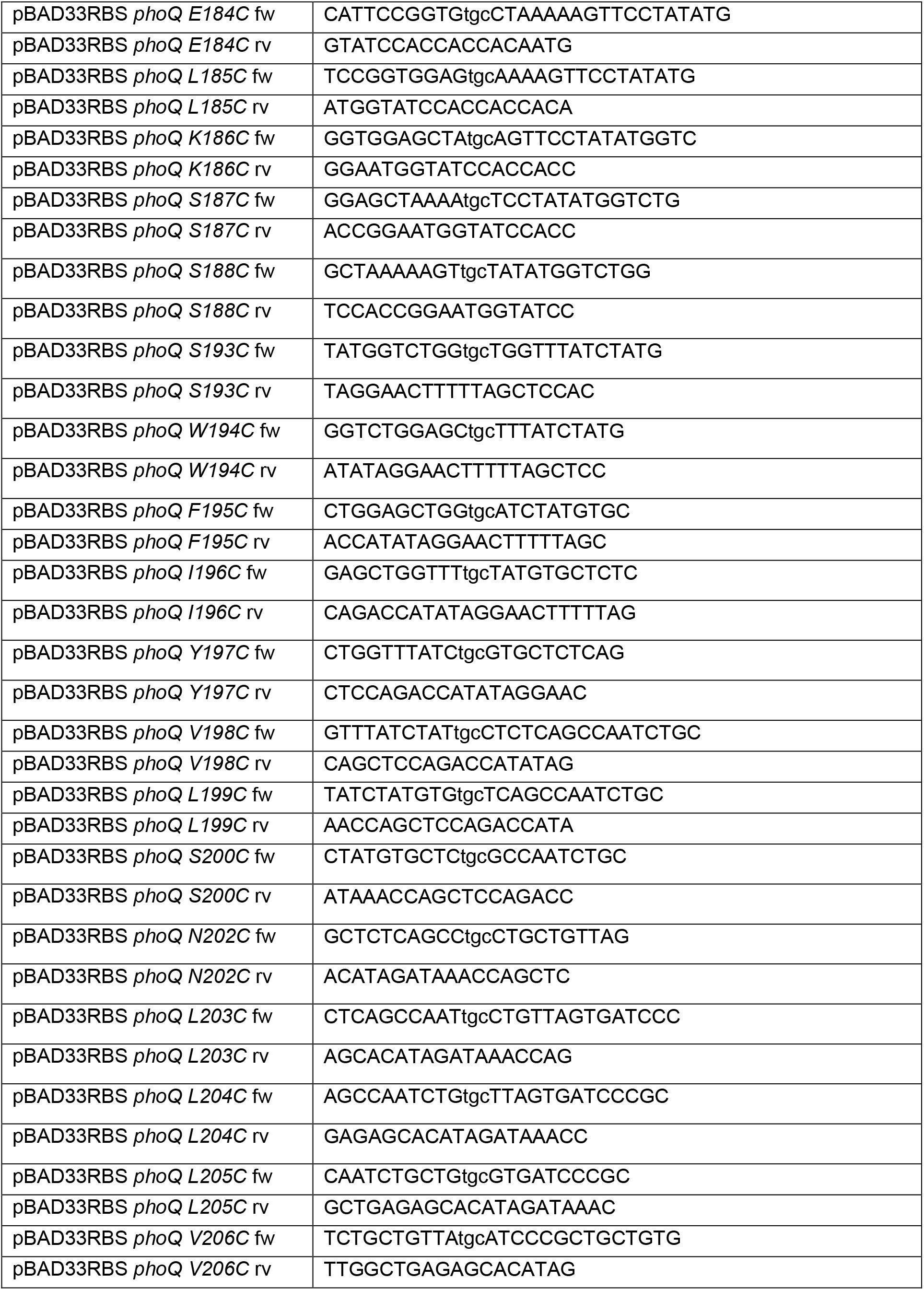
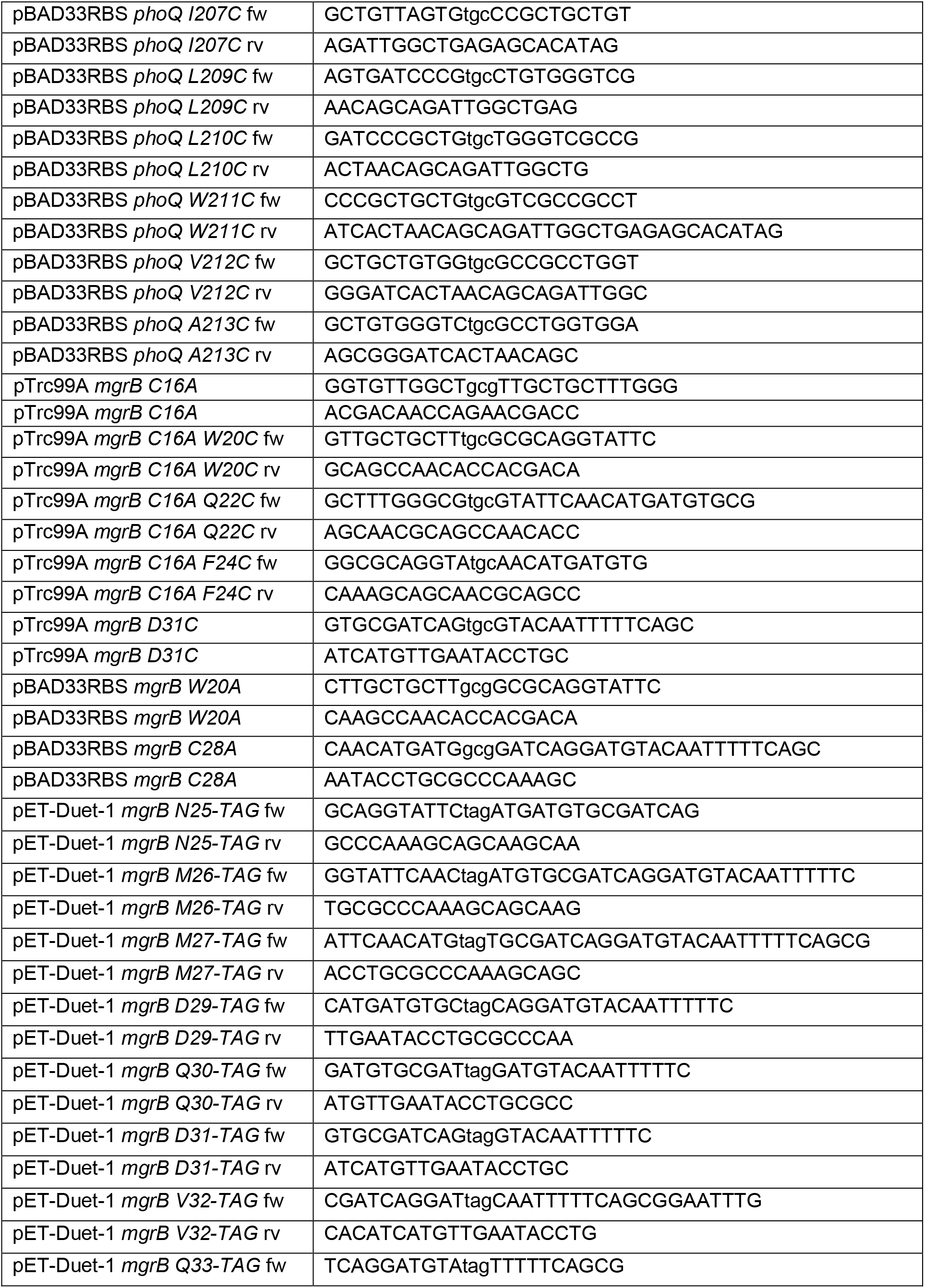
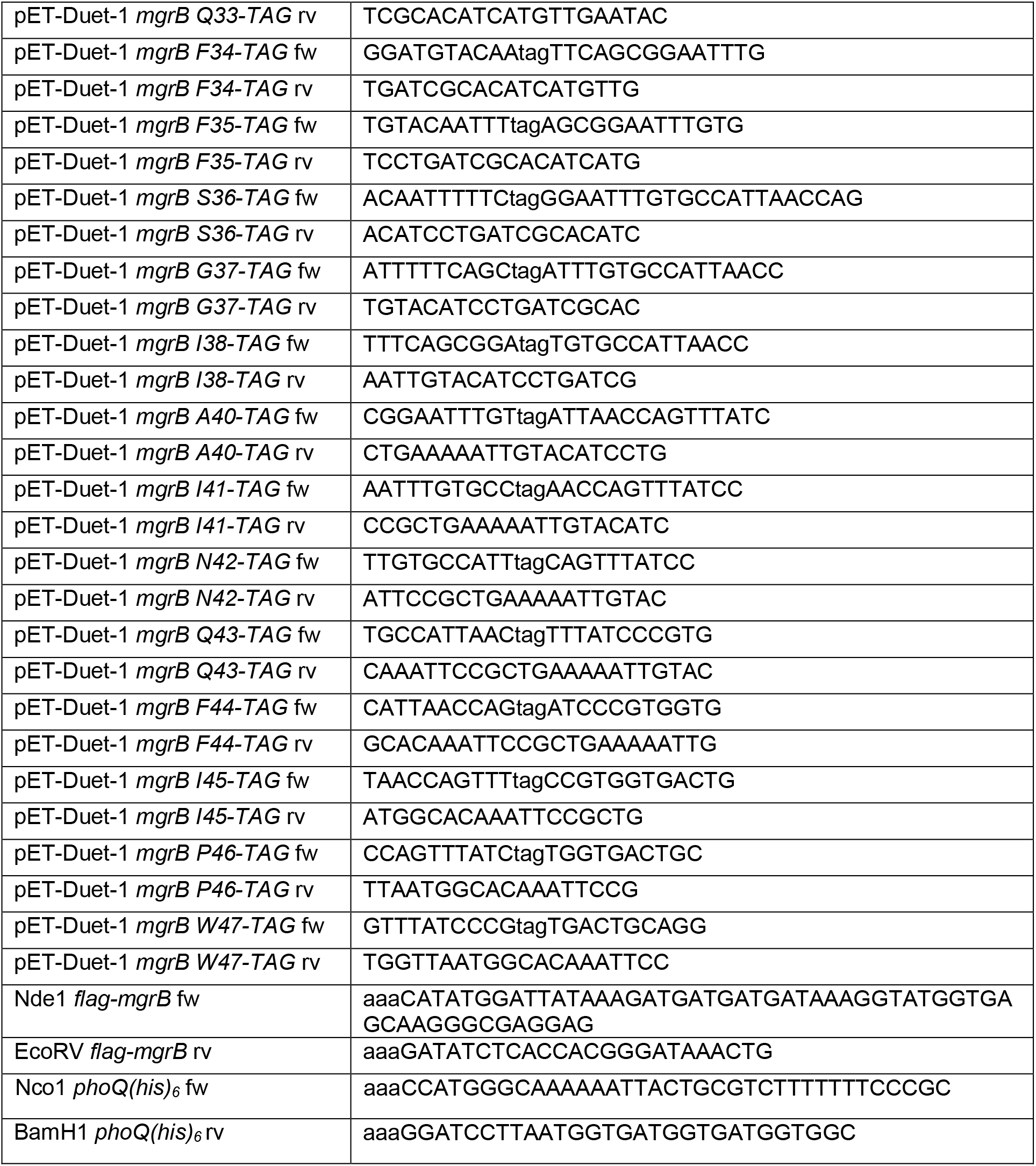
List of primers used in this study.

## References

1. Duval, M. and P. Cossart, Small bacterial and phagic proteins: an updated view on a rapidly moving field. Curr Opin Microbiol, 2017. 39: p. 81–88.

2. Ramamurthi, K.S. and G. Storz, The small protein floodgates are opening; now the functional analysis begins. BMC Biol, 2014. 12: p. 96.

3. Su, M., et al., Small proteins: untapped area of potential biological importance. Front Genet, 2013. 4: p. 286.

4. Weidenbach, K., et al., Small Proteins in Archaea, a Mainly Unexplored World. J Bacteriol, 2022. 204(1): p. e0031321.

5. Sberro, H., et al., Large-Scale Analyses of Human Microbiomes Reveal Thousands of Small, Novel Genes. Cell, 2019. 178(5): p. 1245–1259 e14.

6. Hemm, M.R., J. Weaver, and G. Storz, Escherichia coli Small Proteome. EcoSal Plus, 2020. 9(1).

7. Yadavalli, S.S. and J. Yuan, Bacterial Small Membrane Proteins: the Swiss Army Knife of Regulators at the Lipid Bilayer. J Bacteriol, 2022. 204(1): p. e0034421.

8. Groisman, E.A., A. Duprey, and J. Choi, How the PhoP/PhoQ System Controls Virulence and Mg(2+) Homeostasis: Lessons in Signal Transduction, Pathogenesis, Physiology, and Evolution. Microbiol Mol Biol Rev, 2021. 85(3): p. e0017620.

9. Bader, M.W., et al., Recognition of antimicrobial peptides by a bacterial sensor kinase. Cell, 2005. 122(3): p. 461–72.

10. Prost, L.R., et al., Activation of the bacterial sensor kinase PhoQ by acidic pH. Mol Cell, 2007. 26(2): p. 165–74.

11. Choi, J. and E.A. Groisman, Acidic pH sensing in the bacterial cytoplasm is required for Salmonella virulence. Mol Microbiol, 2016. 101(6): p. 1024–38.

12. Yuan, J., et al., Osmosensing by the bacterial PhoQ/PhoP two-component system. Proc Natl Acad Sci U S A, 2017. 114(50): p. E10792–E10798.

13. Carabajal, M.A., et al., PhoQ is an unsaturated fatty acid receptor that fine-tunes Salmonella pathogenic traits. Sci Signal, 2020. 13(628).

14. Garcia Vescovi, E., F.C. Soncini, and E.A. Groisman, Mg^2+^ as an extracellular signal: environmental regulation of Salmonella virulence. Cell, 1996. 84(1): p. 165–74.

15. Lippa, A.M. and M. Goulian, Feedback inhibition in the PhoQ/PhoP signaling system by a membrane peptide. PLoS Genet, 2009. 5(12): p. e1000788.

16. Eguchi, Y., et al., B1500, a small membrane protein, connects the two-component systems EvgS/EvgA and PhoQ/PhoP in Escherichia coli. Proceedings of the National Academy of Sciences of the United States of America, 2007. 104(47): p. 18712–18717.

17. Yadavalli, S.S., et al., Antimicrobial peptides trigger a division block in Escherichia coli through stimulation of a signalling system. Nat Commun, 2016. 7: p. 12340.

18. Eguchi, Y., et al., Regulation of acid resistance by connectors of two-component signal transduction systems in Escherichia coli. J Bacteriol, 2011. 193(5): p. 1222–8.

19. Salvail, H., J. Choi, and E.A. Groisman, Differential synthesis of novel small protein times Salmonella virulence program. PLoS Genet, 2022. 18(3): p. e1010074.

20. Sanowar, S., A. Martel, and H.L. Moual, Mutational analysis of the residue at position 48 in the Salmonella enterica Serovar Typhimurium PhoQ sensor kinase. J Bacteriol, 2003. 185(6): p. 1935–41.

21. Regelmann, A.G., et al., Mutational analysis of the Escherichia coli PhoQ sensor kinase: differences with the Salmonella enterica serovar Typhimurium PhoQ protein and in the mechanism of Mg2+ and Ca2+ sensing. J Bacteriol, 2002. 184(19): p. 5468–78.

22. Minagawa, S., et al., Isolation and molecular characterization of the locked-on mutant of Mg2+ sensor PhoQ in Escherichia coli. Biosci Biotechnol Biochem, 2005. 69(7): p. 1281–7.

23. Cho, U.S., et al., Metal bridges between the PhoQ sensor domain and the membrane regulate transmembrane signaling. J Mol Biol, 2006. 356(5): p. 1193–206.

24. Cheung, J., et al., Crystal structure of a functional dimer of the PhoQ sensor domain. J Biol Chem, 2008. 283(20): p. 13762–70.

25. Waldburger, C.D. and R.T. Sauer, Signal detection by the PhoQ sensor-transmitter. Characterization of the sensor domain and a response-impaired mutant that identifies ligand-binding determinants. J Biol Chem, 1996. 271(43): p. 26630–6.

26. Venturini, E.S., S. L.; Maaß, S.; Gelhausen, R.; Eggenhofer, F.; Li, L.; Cain, A. K.; Parkhill, J.; Becher, D.; Backofen, R.; Barquist, L.; Sharma, C. M.; Westermann, A. J.; Vogel, J., A global data-driven census of Salmonella small proteins and their potential functions in bacterial virulence. microLife, 2020. 1(1).

27. Shein, A.M.S., et al., High prevalence of mgrB-mediated colistin resistance among carbapenem-resistant Klebsiella pneumoniae is associated with biofilm formation, and can be overcome by colistin-EDTA combination therapy. Sci Rep, 2022. 12(1): p. 12939.

28. Salazar, M.E., A.I. Podgornaia, and M.T. Laub, The small membrane protein MgrB regulates PhoQ bifunctionality to control PhoP target gene expression dynamics. Mol Microbiol, 2016. 102(3): p. 430–445.

29. Lippa, A.M. and M. Goulian, Perturbation of the oxidizing environment of the periplasm stimulates the PhoQ/PhoP system in Escherichia coli. J Bacteriol, 2012. 194(6): p. 1457–63.

30. Yadavalli, S.S., et al., Functional determinants of a small protein controlling a broadly conserved bacterial sensor kinase. J Bacteriol, 2020.

31. Mirdita, M., et al., ColabFold: making protein folding accessible to all. Nat Methods, 2022. 19(6): p. 679–682.

32. Evans, R., et al., Protein complex prediction with AlphaFold-Multimer. bioRxiv, 2022: p. 2021.10.04.463034.

33. Akdel, M., et al., A structural biology community assessment of AlphaFold2 applications. Nat Struct Mol Biol, 2022.

34. Ferris, H.U., et al., Mechanism of regulation of receptor histidine kinases. Structure, 2012. 20(1): p. 56–66.

35. Ryu, Y. and P.G. Schultz, Efficient incorporation of unnatural amino acids into proteins in Escherichia coli. Nat Methods, 2006. 3(4): p. 263–5.

36. Molnar, K.S., et al., Cys-scanning disulfide crosslinking and bayesian modeling probe the transmembrane signaling mechanism of the histidine kinase, PhoQ. Structure, 2014. 22(9): p. 1239–1251.

37. Miller, J.H., A Short Course in Bacterial Genetics - A Laboratory Manual and Handbook for Escherichia coli and Related Bacteria. 1992: Cold Spring Harbor Laboratory Press.

38. Sochacki, K.A., et al., Real-time attack on single Escherichia coli cells by the human antimicrobial peptide LL-37. Proc Natl Acad Sci U S A, 2011. 108(16): p. E77–81.

39. Brink, K.R., et al., An E. coli display method for characterization of peptide-sensor kinase interactions. Nat Chem Biol, 2022.

40. Chamnongpol, S., M. Cromie, and E.A. Groisman, Mg2+ sensing by the Mg2+ sensor PhoQ of Salmonella enterica. J Mol Biol, 2003. 325(4): p. 795–807.

41. Eguchi, Y., et al., The connector SafA interacts with the multi-sensing domain of PhoQ in Escherichia coli. Mol Microbiol, 2012. 85(2): p. 299–313.

42. Yoshitani, K., et al., Identification of an internal cavity in the PhoQ sensor domain for PhoQ activity and SafA-mediated control. Biosci Biotechnol Biochem, 2019. 83(4): p. 684–694.

43. Cabezudo, I., et al., Effect-Directed Synthesis of PhoP/PhoQ Inhibitors to Regulate Salmonella Virulence. J Agric Food Chem, 2022. 70(22): p. 6755–6763.

44. Stopp, M., et al., Transmembrane signaling and cytoplasmic signal conversion by dimeric transmembrane helix 2 and a linker domain of the DcuS sensor kinase. J Biol Chem, 2021. 296: p. 100148.

45. Kuhn, P., et al., Ribosome binding induces repositioning of the signal recognition particle receptor on the translocon. J Cell Biol, 2015. 211(1): p. 91–104.

46. Jo, S., et al., CHARMM-GUI: a web-based graphical user interface for CHARMM. J Comput Chem, 2008. 29(11): p. 1859–65.

47. Best, R.B., et al., Optimization of the additive CHARMM all-atom protein force field targeting improved sampling of the backbone phi, psi and side-chain chi(1) and chi(2) dihedral angles. J Chem Theory Comput, 2012. 8(9): p. 3257–3273.

48. Jorgensen, W.L., Chandrasekhar, J., and Madura, J. D., Comparison of simple potential functions for simulating liquid water. J Chem Phys, 1983. 79(2).

49. Eastman, P., et al., OpenMM 7: Rapid development of high performance algorithms for molecular dynamics. PLoS Comput Biol, 2017. 13(7): p. e1005659.

50. Chow, K.a.F., D. M., Isothermal-isobaric molecular dynamics simulations with Monte Carlo volume sampling. Comp. Phys. Comm., 1995. 91(1-3): p. 283–289.

51. Balusek, C., et al., Accelerating Membrane Simulations with Hydrogen Mass Repartitioning. J Chem Theory Comput, 2019. 15(8): p. 4673–4686.

## SI References

1. Yadavalli, S.S., et al., Functional determinants of a small protein controlling a broadly conserved bacterial sensor kinase. J Bacteriol, 2020.

2. Yuan, J., et al., Osmosensing by the bacterial PhoQ/PhoP two-component system. Proc Natl Acad Sci U S A, 2017. 114(50): p. E10792–E10798.

